# Direct Membrane Penetration of Oligoarginines by Fluorescence and Cryo-electron Microscopy Combined with Molecular Simulations

**DOI:** 10.64898/2026.04.07.716952

**Authors:** Katarína L. Baxová, Mattia I. Morandi, Nadav Scher, Patrik Kula, Ondrej Tichacek, Itay Schachter, Petro Busko, Jiri Zahradnik, Mario Vazdar, Jovi Koikkara, Christoph Allolio, Ori Avinoam, Pavel Jungwirth

**Author notes:** These authors contributed equally to this work (Katarína L. Baxová, Mattia I. Morandi).

## Abstract

Arginine-rich peptides are short amino acid chains capable of spontaneously crossing cellular membranes, with great potential for drug or other cargo delivery. Yet, the mechanisms underlying their cellular penetration are not fully understood. Here, we investigate the modes of action of nonaarginine (R_9_) across membranes of increasing compositional and biological complexity. We combine computational, fluorescence microscopy, and cryo-EM approaches to both visualize the membrane structural changes arising from peptide-lipid interactions and provide a molecular rationale for the observed effects. In large unilamellar vesicles, R_9_ binds preferentially to anionic and PE-rich membranes, induces lipid reorganization, and drives pronounced remodelling, including budding, bifurcations, and time-dependent formation of multilamellar stacks. In cell-derived extracellular vesicles, R_9_-induced remodelling is largely confined to bilamellar bifurcations. In live cells, fluorescent R_9_ forms surface puncta that precede cytosolic entry. Correlative cryo-fluorescence and electron tomography reveals that these puncta correspond to strongly folded, multilamellar membrane structures. We propose that these seemingly contrasting observations can be reconciled within a single R_9_ mechanism of action, involving membrane folding and stacking, where the different observed morphologies arise from the size of the accessible membrane reservoir.

## Introduction

Cell-penetrating peptides (CPPs) are short positively charged chains, typically ranging from 5 to 30 amino acids, with the remarkable ability to traverse cellular membranes.^1^ They achieve this process either through passive translocation across the plasma membrane or via endocytic pathways, enabling the delivery of otherwise membrane-impermeable cargos into cells.^2^ This unique property has sparked significant interest for its potential applications in drug delivery and therapeutic development. Among CPPs, arginine-rich peptides, such as the nonaarginine (R_9_) and the HIV-derived TAT peptide, stand out for their efficiency.^3^

Arginine-rich CPPs have been hypothesized to passively enter cells by forming pores in the membrane^4^ in a certain analogy but also in contrast to antimicrobial pore-forming peptides forming stable pores.^5^ These antimicrobial peptides, whose mechanisms of membrane disruption and self-assembly are reasonably well understood,^6^ provide a stark contrast to arginine-rich CPPs, where the entry process—both in model systems and living cells—remains poorly defined. Recent computational and experimental studies have challenged the pore-formation model for arginine-rich CPPs, suggesting instead that their entry involves more intricate membrane rearrangements and remodelling.^7,8^ Additionally, multiple cell penetration mechanisms can operate simultaneously, including both active (involving ATP) and passive processes. Recently, we proposed^9^ that CPPs can passively cross membranes by inducing multilamellarity, a phenomenon observed in lipid model systems across multiple studies where arginine-rich peptides interact with membranes. The proposed entry mechanism involves the membrane wrapping that leads to a pore formation. In addition to the complexity of the problem, the high levels of peptide-induced aggregation, adhesion, and fusion observed in experimental model systems obscure the elucidation of the exact peptide’s mode of action on membranes.

To fill this knowledge gap, alongside with addressing the challenges in translating findings on lipid model systems to cellular membranes, we dissected the problem into questions how nonaarginine remodels membranes of increasing compositional complexity, ranging from simple composition liposomes in the form of large unilamellar vesicles (LUVs) to actual cellular membrane compositions, both in the form of extracellular vesicles (EVs) and live cells. Using a combination of molecular modelling, cryogenic transmission electron microscopy (cryo-TEM), cryo-electron tomography, live cell imaging, and correlative light and electron microscopy (CLEM), we demonstrate that arginine peptides remodel membranes even in the absence of inter-vesicular interaction or aggregation. Multilamellarity is observed when there is a sufficient amount of extra lipids, as is the case for cells or interacting vesicles. We rationalize our findings in terms of a combination of the principal peptide action on membranes, i.e., folding by lipid re-arrangement and bilayer stacking, and the presence or absence of an available membrane reservoir.

## Materials and Methods

### Umbrella Sampling Simulations

All molecular dynamic simulations were performed in the Gromacs 2022 package using CHARMM-GUI^10^ to build lipid bilayers.

The isothermal-isobaric ensemble was set at 303.15 K by Nose-Hoover temperature coupling^11^ for the groups of protein, membrane, and water with ions with a time constant of 1 ps and semiisotropic Parrinello-Rahman pressure coupling^12^ at 1 bar. Particle-Mesh Ewald electrostatics (PME) was used to treat long-range interactions with a 1.2 nm cutoff.^13^ Bonds with hydrogen atoms were constrained by LINCS,^14^ and the SETTLE algorithm was used for constraints in water molecules.^15^ The leap-frog integration algorithm timestep was set to 2 fs.

In order to avoid one of the common flaws of current non-polarisable force-fields, excessive binding of (biologically relevant) charged species,^16^ a charge scaling CHARMM36m/CHARMM36 force field derivative - prosECCo75^17^ was used alongside the traditional CHARMM36/LJ-PME force field with the TIP3Ps water model.^18^

In order to obtain the Potential of Mean Force (PMF) of a peptide-membrane interaction, we employed Umbrella sampling simulations with umbrella windows constructed along the membrane-peptide center of mass (COM) in the z direction. Each PMF profile was constructed using at least 18 umbrella windows (corresponding to at least 3.6 *µ*s of simulation time per single PMF profile) with membrane-peptide distances and their corresponding force constants recorded in Table S1.^19^ Each window was simulated for 200 ns with at least 10 ns preequilibration.

The simulated systems contained 128 lipids per leaflet, 100 waters per lipid, one peptide -tetraarginine (R_4_), nonaarginine (R_9_) or nonalysine (K_9_). Potassium and chloride ions were added to mimic the biological concentration of 150 mM and, moreover, to balance the overall system charge to 0, since the investigated peptides are positively charged.

Due to the high computational cost of the Umbrella sampling simulation, only the Pros-ECCo75^17^ force field parameters were used in this case, except for the R_9_ with DOPE:DOPC:DOPS:chol 40:20:20:20 membrane composition case, which was simulated also using regular CHARMM force field for comparison (see SI Fig. S3).

### Chemicals

Lipids palmitoyl-oleoyl-glycero-phosphocholine (POPC), dioleoyl-glycero-phosphoethanolamine (DOPE), dioleoyl-glycero-phospho-L-serine (DOPS), dioleoyl-glycero-phosphocholine (DOPC), and cholesterol were purchased from Avanti Polar Lipids, Inc. (Alabaster, USA).

Nonaarginine (R_9_), tetraarginine (R_4_) and nonalysine (K_9_) peptides, as well as their Tamra-labelled forms, were custom-made in-house at IOCB, using the procedure described in Ref.^20^

### Large Unilamellar Vesicle Preparation

Large Unilamellar Vesicles (LUVs) were prepared similarly to,^9^ with lipid compositions (molar ratios) of POPC, DOPE:DOPC:DOPS 60:20:20 (DDD), or DOPE:DOPC:DOPS:chol 40:20:20:20 (DDDC). Briefly, lipids dissolved in chloroform were mixed and left to evaporate overnight under vacuum. The lipid film formed this way was rehydrated in PBS -/- buffer (or regular PBS +/+ buffer, see Supporting Information), pre-warmed to room temperature. After vortexing, the lipid solutions were placed in a 37^◦^C water heat bath for 1 h. After another vortexing, subsequent 10 min sonication, and final vortexing, lipid suspensions were extruded 21x through a 100 nm pore size polycarbonate membranes (Avanti Polar Lipids).

The LUVs were stored at 4^◦^C and used within 2 weeks after preparation.

### Laurdan Kinetics Measurements

At the beginning of the LUV preparation, the amount of Laurdan in DMSO, corresponding to a laurdan:lipid molar ratio of 1:100, was added to the initial lipids in chloroform solution. The LUVs were dissolved in PBS buffer to a final concentration of 0.4 mM, stirred, pre-warmed, and kept at 20^◦^C by a water bath during measurements. The amount of peptide corresponding to the peptide:lipid ratio of 1:1 was added to the LUV suspension during the measurements.

Measurements were obtained using a Varian Cary Eclipse Fluorescence Spectrophotometer (Agilent). Laurdan fluorescence emission, excited at 350 nm, was measured at 440 nm (*I*_440_) and 490 nm (*I*_490_) each 0.5 s to obtain the general polarization factor (*GP* ), defined as *GP* = (*I*_440_ − *I*_490_)*/*(*I*_440_ + *I*_490_).^21^ The ΔGP is calculated as a difference between the average GP during the three-minute time interval before the addition of peptide (or buffer control) and the average GP during 2 − 5 min after the peptide addition.

Freshly prepared EVs were diluted 10 times in PBS -/- and consequently incubated with Laurdan to its final concentration of 5 *µ*M in a 37^◦^C water bath for 30 min. For each measurement, 200 *µ*l of the EV-laurdan solution was dissolved in PBS -/- buffer to a total volume of 500 *µ*l. The samples were preheated and measured at 37^◦^C. 40 *µ*l of 10 mM peptide was added during kinetics measurements to obtain the GP factor.

### Cell Culture

Human bone osteosarcoma (U-2 OS, ATCC HTB-96™) cells were cultured under standard conditions in DMEM media (Thermofisher Scientific) supplemented with 10% fetal bovine serum (FBS, Thermofisher Scientific), 1% penicillin-streptomycin mixture, and 1% glutamine solution, in an incubator at 5% CO_2_ and 37^◦^C. These cells were used for live cell confocal imaging, flow cytometry, and CLEM experiments.

The ovarian cancer OVCAR-3 cell line was used to produce the extracellular vesicles (EVs). 10x10^6^ cells were seeded in a 175 cm^2^ flask in culture media composed of DMEM with 10% EV-free FBS, 1% sodium pyruvate, 1% l-glutamate, and 1% Pen/Strep.

### Extracellular Vesicle Isolation

Extracellular vesicles (EVs) derived from the ovarian cancer cells (OVCAR-3; ATCC-HTB-161) conditioned media were harvested as previously described.^22^ Upon reaching ∼ 70% confluency (typically 2 days post-seeding), cells were washed twice with PBS buffer without Ca^2+^ and Mg^2+^ (PBS -/-) and replenished with naive EV-free growth medium. Cell culture media was collected after 48 h and spun at 300 g for 10 min at 4^◦^C to remove large debris and remaining cells. The supernatant was collected and spun at 2000 g for 10 min at 4^◦^C. The supernatant was then collected, spun at 10000 g for 45 min to remove larger vesicular particles, and filtered through a 0.22 *µ*m polycarbonate filter. The resulting media was used for vesicle isolation within 2 days or frozen at −80^◦^C for biochemical analysis. Following filtration, the cell culture media was spun using an ultracentrifuge, a Ti45 rotor (Beckman Coulter, Fullerton, CA, USA) at 100000 g for 4 h at 4^◦^C. The supernatant was removed, and the resulting pellet was washed once with PBS -/-, and then resuspended in PBS -/-.

Following differential ultracentrifugation, EVs were fractionated by OptiPrep density gradient ultracentrifugation (100000 g, 18 h, 4^◦^C) using a SW41 rotor (Beckman Coulter, Fullerton, CA, USA) through a continuous 5% to 40% OptiPrep (Sigma-Aldrich, D1556) gradient. The fractions (1 ml) were collected from the top of the gradient for further analysis, and the density was verified by measuring the mass of a 100 *µ*l aliquot of each fraction. The EV-specific density fractions were then pooled together and subsequently concentrated via ultracentrifugation (100000 g, 4 h, 4^◦^C) through a 20% w/v sucrose cushion in a SW41 rotor (Beckman Coulter, Fullerton, CA, USA). The resulting supernatant was discarded, and the EV pellet was resuspended in PBS -/-.

### Preparation of Vesicle-depleted Fetal Bovine Serum (EV-free FBS)

FBS was depleted from extracellular vesicles by two rounds of ultracentrifugation at 100000 g for 18 h in a Beckman Ti45 rotor. In each round, the supernatant was collected, and the large pellet at the bottom of the tube was discarded. After the final round of ultracentrifugation, the supernatant was collected and filtered through a 0.22 *µ*m pore membrane, aliquoted, and stored at −20^◦^C for the preparation of an EV-free growth medium.

### Confocal Live Cell Imaging

U-2 OS cells were seeded into a 35 mm ibidi µ-Dish (approximately 300000 cells per dish) and grown overnight. After a PBS wash, cells were incubated in 1:2000 Hoechst-containing pre-warmed PBS media for 10 min and subsequently washed 3x with PBS. If using CellMask Green, cells were incubated with freshly made 1:1000 staining solution for 10 min and washed 3x with PBS.

During the cell imaging, cells were incubated in Hank’s Balanced Salt Solution buffer containing Ca^2+^ and Mg^2+^ ions (HBSS +/+, Gibco). After checking the cell viability in a differential interference contrast (DIC) channel (30 ms exposure), HBSS buffer was gently aspirated from the dish, and a pre-warmed Tamra-labelled peptide, diluted in HBSS, was gently added. The cells were then examined by a confocal microscope (inverted Olympus IX83, equipped with OKOLab Microscope Incubator) at 37^◦^C and under 5% CO_2_ atmosphere. 40x NA 1.3, or 100x NA 1.49 oil immersion objectives were used. VisiView software was used to control the image acquisition, and ImageJ/Fiji for processing. All images shown have been adjusted for brightness and contrast.

The Hoechst dye for nuclei labelling was visualised in a widefield channel, excited at 405 nm with an exposure time of 15 ms and a laser power of 5 mW. The 561 nm confocal channel showing Tamra-labelled peptides was used with a laser power of 15 mW and 120 ms exposure. The signal of CellMask Green, a membrane-labelling dye, was collected by a confocal 488 nm laser channel with 120 ms exposure and 15 mW laser power.

To obtain cell-penetration statistics after 20 min peptide incubation, image sets were collected with two different magnifications. The images acquired with 100x magnification were stitched and smoothened to remove stitching artefacts with the BaSiC plugin.^23^ No significant changes were observed compared to results without stitching smoothening. In the case of larger fields of view acquired using 40x magnification, an average intensity of up to the 3 sharpest *z*-planes was used for further processing. For each individual image, at least 3 different regions outside the cell boundaries were selected and the average intensity of the Tamra signal was calculated, serving as a reference. For each living cell, the Hoechst channel was used to determine the boundary of the nucleus region. Since every time the peptide was observed to translocate into the cell, it also promptly reached the cell nucleus, we can compare the calculated average Tamra-peptide intensity inside the cell nucleus region to the outside reference. If this average nucleus region intensity was higher, the cell was considered penetrated. The final cell penetration statistics after 20 min peptide incubation were acquired as a weighted average of several biological and technical repeats, including hundreds of cells per condition.

### Flow Cytometry

U-2 OS cells were cultured in the 150 mm dishes to ∼ 70% confluency. For flow cytometry experiments, cells were washed with PBS buffer, trypsinized, resuspended in 10% FBS supplemented DMEM, centrifuged for 5 min at 160 RCF, (media aspired) and finally resuspended in HBSS media.

For peptide concentration measurements, (Figure 7, D), 1 ml of the cell suspension was incubated with 1 *µ*M, 5 *µ*M, 10 *µ*M and 15 *µ*M peptide, or no peptide as a control, for 20 min in the incubator at 5% CO_2_ and 37^◦^C. After 5 min centrifugation (Labnet Prism mini centrifuge), the supernatant was aspirated, cells were placed on ice, and promptly transferred to be resuspended in HBSS and measured immediately on the CytoFLEX Flow Cytometer (Beckman Coulter). The cells were first gated to show live cells only (see Figure S14), then subsequently to exclude multiplets. Fluorescence intensity histograms (channel PE-A, Tamra) were calculated from this gate.

Flow cytometry measurements were evaluated with respect to the elevated fluorescent background associated with unbound fluorescently labelled peptides. When cellular fluorescence was weak, the background signal overlapped with cell-derived fluorescence, resulting in broader intensity distributions and, in some cases, apparent negative values following background correction. A comparable effect was observed in confocal microscopy experiments, where diffuse extracellular fluorescence masked the low intracellular signal.

For time-resolved flow cytometry measurements (Figure 7, E), the peptide was added directly to U-2 OS cells suspended in HBSS buffer, quickly mixed and immediately measured on BD FACSAriaTM Fusion. The samples were agitated at 100 RPM with the temperature maintained at 4^◦^C. The gating strategy and data analysis is shown in Fig. S16. Briefly, 10 min of the measured data were used. First, collected events were sorted into 3 categories: subcellular fragments (small), cells, and aggregates (large). Secondly, singlets were extraxted from the cell population. Lastly, for these selected events, the fluorescence intensity evolution in time was extracted.

Additionally, Tamra-labelled peptides in HBSS +/+ buffer, without the presence of cells, were measured for 5 min.

### Cells CLEM Measurements

For this experiment, Quantifoil R3.5*/*1 200mesh Au grids were plasma-cleaned using a remote plasma cleaning mode (Tergeo plasma cleaner, PIE Scientific). The grids were left in a tissue culture hood under UV for 1 hour before being transferred into a 14 mm glass-bottomed well of a 35 mm imaging plate (D35-14-1.5N, Cellvis). The glass-bottomed well contained 350 *µ*l of pre-heated growth medium, with the Quantifoil layer facing upward. Then, 200 *µ*l of the growth media was replaced with the same media, containing 30000 U-2 OS cells per well, and left to settle for 2-3 hours. After cell adhesion, 1 ml of growth media was gently added, and the cells were incubated overnight at 5% CO_2_ and 37^◦^C.

Right before plunge-freezing, cells were washed with warm PBS and incubated with Hoechst dye (1:2000) for 10 min under 5% CO_2_ and 37^◦^C. The cells were washed three times with warm PBS. The grids were then incubated for 3 minutes with 5 *µ*l HBSS buffer containing 10 *µ*M Tamra-R_9_ at 37^◦^C and 90% humidity in the chamber of an EM-GP plunge freezer (Leica Microsystems). Right before plunging, the grids were back-blotted on an ashless Whatman in the plunger with a blotting time of 2.5-3 s. The grids were then clipped in a clipping station (Thermofisher Scientific) and stored in an autoloader grid boxes in a grid storage system (Subangstrom) until imaged by Cryo fluorescence microscopy (cryo-FM).

Cryo-FM was performed using an EM CryoCLEM cryo-light microscope (Leica Microsystems) as previously described.^24–26^ The grids were transferred to a pre-cooled transfer shuttle of the microscope. The samples were imaged with an HCX PL APO 50X CLEM cryo-air objective and an ORCA-flash 4.0 camera (Hamamatsu Photonics) with DAPI (ex350*/*50; em460*/*50) and TXR (ex560*/*40; em630*/*75) filter cubes. Initially, a spiral scan was acquired in the blue (DAPI) and brightfield channels to roughly estimate the quality of the sample and freezing. Then, a mosaic of good areas was created with a step size of 1-5 *µ*m over 25-50 *µ*m. Lastly, individual tiles were acquired in a 250 nm step size over 10 *µ*m for regions of interest.

Cryoelectron tomography (Cryo-ET) acquisition of selected fluorescent point of interest was performed on a Titan Krios 3Gi STEM/TEM, Thermo Fisher Scientific, USA) at 300 kV located at the EMBL Imaging Center. Medium maps corresponding to individual region-of-interest tiles were acquired at a nominal magnification of 2, 250x, and correlated to cryo-FM tiles using eC-CLEM.^27^ Correlated maps were then used to select tomographic acquisition points. The tilt series were acquired in low-dose mode using SerialEM software at a nominal magnification of 21000x with an angular range from −60^◦^ to +60^◦^, an angular increment of 2^◦^ using a −6.5 *µ*m defocus, 0.671 nm per pixel, and a maximal total dose of 140 e^−^/Å^2^. For all datasets, tomograms were reconstructed using the weighted back-projection technique in the IMOD software suite^28^ with a SIRT-like filter equivalent to three iterations.

### Preparation of Cryo-TEM Samples

Cryo-EM samples of both EVs and LUVs were prepared on either 300-mesh 2*/*2 copper Quantifoil or C-flatm or lacey EM grids (Electron Microscopy Sciences, USA), on which 10 nm colloidal gold particles (Au–NP) were preadsorbed (Aurion, Netherlands). The Au–NP adsorbed grids were then glow-discharged on either a Pelco EasyGlow system (30 s, 25 mA). An aliquot (2.0 *µ*l) of the aqueous solution of the LUV sample was applied to the carbon side of EM grids with subsequently another aliquot of the desired peptide solution or buffer added directly on the grid side, which was then incubated in the humidity chamber of the instrument for the desired time at 100% humidity and room temperature, and subsequently blotted for 2.0 s at blot force −7 and plunge-frozen into the precooled liquid ethane with a Vitrobot Mark IV (Thermo Fisher, USA).

### Cryo-TEM Acquisition

The micrographs of vitrified samples were collected on three separate microscopes: i) a Talos Arctica G3 TEM/STEM (Thermo Fisher Scientific, USA), equipped with a OneView camera (Gatan) at an accelerating voltage of 200 kV, located at the Weizmann Institute of Science, ii) a Jeol JEM-2100PLUS, equipped with a high-tilt Gatan cryo-holder and a TVIPS TemCam-XF416 CMOS camera, located at IOCB, and iii) a Talos Arctica G3 TEM/STEM located at CEITEC. For all three microscope configurations, grid and square mapping, as well as image acquisition, were performed using SerialEM software.^29^ The grid and square mapping were performed at nominal magnification of 180x and 13, 500x, respectively, for configuration (i), and x80 and x2500 for configuration (ii), and x80 and x8700 for configuration (iii). High-magnification images were recorded at 73000x nominal magnification (0.411 nm pixel size) with a −2.5 *µ*m defocus value for configuration (i), 60000x nominal magnification (0.1939 nm pixel size) with a −2.1 defocus value for configuration (ii) and 56000x nominal magnification (0.211 nm pixel size) with a −2.1 defocus value for configuration (iii). To minimize radiation damage during image acquisition, the low-dose mode in SerialEM software was used with the electron dose kept below 100 e^−^/Å^2^.

### Cryo-Electron Tomography

Samples were prepared as for cryo-TEM (described above) with some modifications. Prior to plunging, samples were mixed 50:1 with a suspension of 10 nm Au–NP (Aurion, Netherlands) to serve as fiducial markers for reconstruction. Tilt series were collected using either A) a transmission electron microscope (Titan Krios 3Gi STEM/TEM, Thermo Fisher Scientific, USA) at 300 kV equipped with a Gatan K3 direct detector mounted at the end of a Gatan BioQuantum energy filter set in zero-energy–loss mode (slit width, 20 eV), located at the Weizmann Institute of Science, B) a Titan Krios 3Gi STEM/TEM at 300 kV equipped with a Gatan K3 direct detector mounted at the end of a Gatan BioQuantum energy filter set in zero-energy–loss mode (slit width, 20 eV), located at CEITEC, and C) a Titan Krios 3Gi STEM/TEM, Thermo Fisher Scientific, USA) at 300 kV located at the EMBL Imaging Center.

For each configuration, tilt series were acquired according to the following parameters:

The tilt series were acquired in low-dose mode using SerialEM software at a nominal magnification of 42000x with an angular range from −60^◦^ to +60^◦^, an angular increment of 4^◦^ using a −2.5 *µ*m defocus, 70 *µ*m objective aperture, 0.214 nm per pixel, and a maximal total dose of 120 e^−^/Å^2^.
The tilt series were acquired in low-dose mode using SerialEM software at a nominal magnification of 42000x with an angular range from −60^◦^ to +60^◦^, an angular increment of 3^◦^ using a −2.5 *µ*m defocus, 0.208 nm per pixel, and a maximal total dose of 120 e^−^/Å^2^.
The tilt series were acquired in low-dose mode using SerialEM software at a nominal magnification of 64000x with an angular range from −60^◦^ to +60^◦^, an angular increment of 3^◦^ using a −2.5 *µ*m defocus, 0.197 nm per pixel, and a maximal total dose of 120 e^−^/Å^2^.

For all datasets, tomograms were reconstructed using the weighted back-projection technique in the IMOD software suite with a SIRT-like filter equivalent to three iterations,^28^ following nonlinear anisotropic diffusion (NAD) de-noising^30^ if indicated.

### Machine Learning Analysis of Cryo-ET Data

The tomograms measured by cryo-EM and processed into *z*-stacks were plane-cropped to show only the relevant space containing lipid vesicles. Each cropped tomogram was decomposed into overlapping 64×64 pixel patches using a sliding window with a stride of 2 pixels. A two-stage convolutional neural network (CNN) classification pipeline was used. In the first stage, CNN classifier detected whether a given patch contained vesicle membrane. Only patches identified as membrane-containing were passed to the second stage.

We used a sequential CNN architecture consisting of three blocks of paired 3×3 convolutions (32 to 64 filters) followed by max-pooling.^31^ Regularization was prioritized through integrated random flip augmentation, progressive dropout (*p* = 0.25 to 0.5), and categorical cross-entropy loss with a label smoothing factor of 0.2. The model was trained using the Adam optimizer^32^ over 200 epochs.

The second-stage CNN then classified each patch according to specific membrane properties, such as multilamellarity or the presence of peptides on the lipid membrane (see the example in Figure S12).

Both the membrane detector and the feature classifier were trained on a manually labelled dataset. The Cryo-EM patches were annotated by expert inspection to ensure accurate ground truth. After each training cycle, the resulting prediction masks and labelled regions were manually reviewed. Misclassified regions were isolated to form an additional fine-tuning dataset, specifically enriched with difficult or ambiguous examples. The model was retrained iteratively on this expanded dataset. This procedure was repeated until no further significant improvement in classification performance was observed.

After classification, the number of patches assigned to each morphological class was quantified for each tomogram *z* slide. Then, they were summed across the whole tomogram *z*-stack belonging to an individual LUV vesicle. A fraction of these morphological class sums was used as a global descriptor of the said vesicle.

### Statistical Analysis of Membrane Remodelling Uniformity

Membrane remodelling frequencies were quantified per cell from cryoEM images as the proportion of points exhibiting multilamellar morphology out of total points imaged per cell (*n* = 58 points across 18 cells). Variability across cells was assessed using a strip plot with 95% Wilson score confidence intervals, which account for small sample sizes (*n* = 1– 11 points/cell) via continuity correction and center extremes toward the population mean. Wilson CIs were calculated for each cell using the formula:

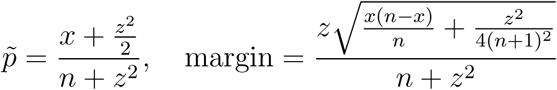

where *x* = remodelling successes, *n* = points imaged, *z* = 1.96 (95% CI), yielding lower/upper bounds 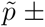 margin. Point sizes were scaled by *n* per cell; cells were sorted by ascending frequency. Uniformity was confirmed by all CIs overlapping the global proportion (27*/*58 = 46.6%) and a non-significant Kruskal–Wallis test (*H* = 17.00, *p* = 0.4544). Analysis and plotting were performed in OriginPro 2026 using pre-calculated Wilson CI columns.

## Results and Discussion

The mammalian membrane is a complex system containing various lipids and proteins with different compositions at the inner and outer cell membrane leaflets. In addition to live-cell experiments, where the (human bone osteosarcoma) U-2 OS cell line was chosen for its flat morphology, suitable for cryo-ET acquisitions without thinning down the sample, we selected the following simplified systems for molecular simulations of membranes, fluorescence spectroscopy, and cryo-EM measurements on LUV vesicles. First, the POPC-only lipid composition mimics the outer leaflet of the mammalian cell membrane, characterized by its charge neutrality and a rich phosphatidylcholine (PC) content. Second, the lipid composition of DDD serves as a simple representation of the inner membrane leaflet, mimicking its negative charge. Third, in order to investigate the effect of cholesterol, another lipid composition,DDDC, was also considered.

### Peptide-Membrane Binding Strength by Umbrella Sampling Simulations

The free energy profiles of the peptide-membrane interactions for R_9_, K_9_, and R_4_ for the three different lipid compositions are presented in Figure 1, with calculated free energy minima positions and the corresponding free energy values given in Table 1.

**Figure 1:**
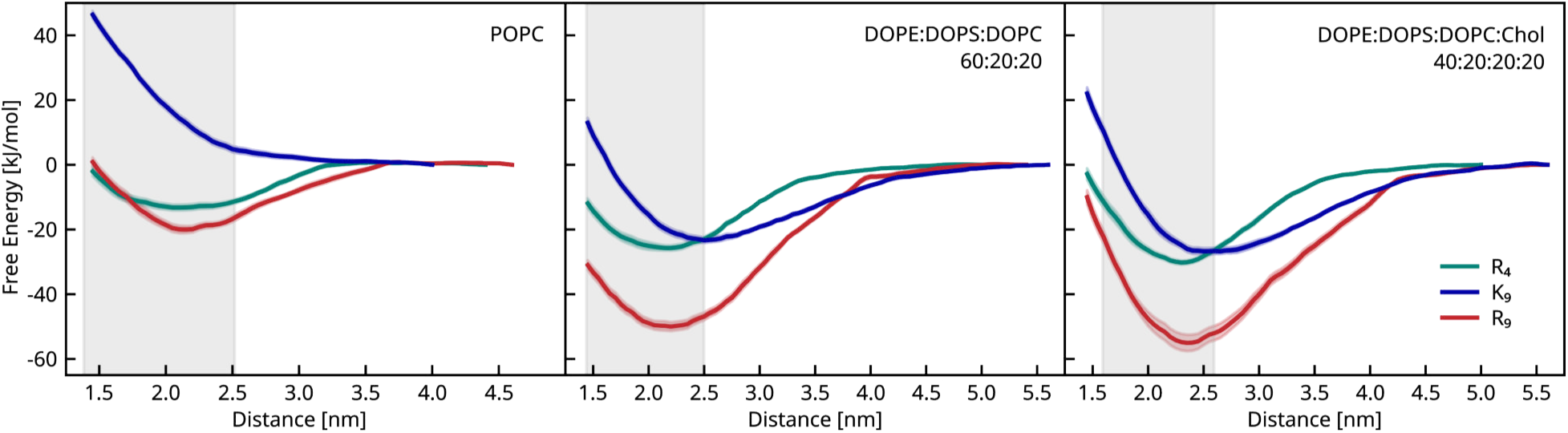
Free energy profiles as functions of the distance between the peptide and membrane centers of mass. POPC (left), DDD (middle), and DDDC (right) membrane compositions. Peptides are colored as follows: R_9_ (red), K_9_ (blue), and R_4_ (green). The gray area marks the phosphate-rich membrane region.

**Table 1:** Distance positions (*d*) and energies (*E*) of the calculated free energy profiles minima depicted in Figure 1.

|  | POPC |  | DDD |  | DDDC |  |
| --- | --- | --- | --- | --- | --- | --- |
| | $d$ [nm] | $E$ [kJ/mol] | $d$ [nm] | $E$ [kJ/mol] | $d$ [nm] | $E$ [kJ/mol] |
| $R_4$ | 2.10 | $-13.2 \pm 0.9$ | 2.15 | $-25.7 \pm 0.8$ | 2.30 | $-30.2 \pm 0.6$ |
| $R_9$ | 2.15 | $-20.0 \pm 1.1$ | 2.20 | $-50.0 \pm 1.4$ | 2.35 | $-55.0 \pm 2.6$ |
| $K_9$ | <sup>a</sup> | <sup>b</sup> | 2.50 | $-23.3 \pm 0.8$ | 2.65 | $-26.7 \pm 0.9$ |

<sup>a</sup>The POPC- $K_9$ profile does not have a close-distance minimum, but rather converges to zero at large ("infinite") distance,
<sup>b</sup>therefore, the value of binding free energy can be considered zero.

In case of a neutral membrane containing only zwitterionic POPC lipids (Figure 1, left), there is no attraction between the membrane and nonalysine.^33,34^ In contrast, the minima corresponding to the interaction strengths of −13.2 kJ*/*mol and −20.0 kJ*/*mol (shown in Table 1) are observed for R_4_ and R_9_, respectively, suggesting that the chemical specificity of the cationic side chain group, as well as the peptide length, play deciding roles in the peptide-POPC bilayer interactions.

Figure 1 middle and right, shows peptide interactions with mixed negatively charged membranes with and without cholesterol. In both cases, all three peptides prefer close contact with membranes. Tetraarginine and nonalysine interact with membranes with a comparable strength - in the case of DDD membrane, the free energy minimum is 25.7 kJ*/*mol and 23.3 kJ*/*mol for R_4_ and K_9_, respectively. Similarly, for the cholesterol-containing membrane composition DDDC, the free energy minimum is 30.2 kJ*/*mol and 26.7 kJ*/*mol, respectively. Nonaarginine attaches approximately twice as strongly as the other two peptides by - 50.0 kJ*/*mol or 55.0 kJ*/*mol for membranes without or with cholesterol, as seen in Table 1. In general, cholesterol presence in the membrane causes slightly deeper interaction minima for all peptides (Figure 1, right), compared to the membrane without cholesterol (Figure 1, middle). For all membrane-lipid systems investigated, both polyarginines penetrate deeper into the membranes than nonalysine, as seen in Figure 1 and Table 1.

Throughout simulations, negative DOPS lipids prefer to stay in a close contact with positively charged peptides, as can be seen in the DDD system density maps (Figure S1). This occurs mainly at the expense of DOPC lipid density, while DOPE lipid density remained only slightly altered in the windows exhibiting peptide-membrane proximity. Similarly, in the case of the cholesterol-containing membrane system (Figure S2), negative DOPS is the preferred counterpart to positively charged peptides, with DOPC lipids retreating from their vicinity. The cholesterol density increases in the membrane region close to the R_4_ or K_9_, but not the R_9_ peptide.

To compare the CHARMM force field with the charge-scaled ProsECCO, the free energy profile of R_9_ interacting with the DDDC membrane was calculated also using CHARMM (see Figure S3), resulting in a much stronger interaction.

### Continuum Elastic Models

#### I. Monte Carlo Simulations

In the simulations presented in the previous section, lipid membranes were kept flat by employing periodic boundary conditions. Nevertheless, earlier studies^9,35,36^ have indicated that peptide binding may also influence membrane curvature.

In recent work, we constructed and validated a continuum elastic model for the binding and curvature generation of R_9_.^37^ The model departs from detailed simulations of R_9_ and K_9_ adsorbing on lipid bilayers and was parametrized using the ReSIS method of extracting bending rigidities^38^ and a local stress code.^39^ The parameters reflect that R_9_ (and to a lesser extent K_9_) induce negative curvature on DOPS and also on DOPE lipid membranes. The model uses a Helfrich Hamiltonian,^40^ with coefficients depending on the local lipid composition. It includes ideal lipid mixing, lipid-specific protein binding, area compressibility, and volume restricted by osmotic pressure. The published results were obtained by Monte-Carlo simulation of a dynamically triangulated mesh within the OrganL software.^41^ The large size of the peptide with respect to the lipids (covering 21-23 lipids at a time), together with the large binding energy to, in particular, DOPS, allows it to overcome the mixing entropy and induce curvature sorting of the lipids together with R_9_. This sorting of proteins occurs in concomitant with mean curvature, as well as negative Gaussian curvature, i.e., in the bud and the neck of the invagination. Our results, summarized graphically in Figure 2 A, show that the R_9_ curvature induction is by itself capable of generating stable invaginations even in small vesicles. The occurrence of an invaginated geometry in a closed vesicle leads to a sphere-in-a-sphere geometry, typically called a stomatocyte. The stomatocyte geometry is stabilized by the presence of R_9_, (but not K_9_) over the spheroid shape found for a CPP-free vesicle. We found that the stomatocyte is stable under the condition that, e.g., a 100 nm diameter vesicle is filled to no more than about 85%. This means that a certain amount of excess membrane area over volume is required to stabilize this structure. In a small vesicle, the osmotic pressure rapidly increases with the effective concentration of salt as a result of liposome invagination. This illustrates how much easier such deformations are to achieve, e.g., on GUVs or rough cellular surfaces. In its published version, the model did not include any peptide-peptide interactions beyond size exclusion. Accordingly, our initial model^37^ neglected the evident contribution of crosslinking via R_9_. Crosslinking can be expected to stabilize tight, long necks in a similar way as it stabilizes multilamellar structures. As such structures were observed in the data, we have included through-space intermembrane interactions in the form of a Lennard-Jones type R_9_-R_9_ interaction. This effective potential only applies on opposing faces containing R_9_ molecules. We chose repulsion such that the minimum of the potential aligns approximately with the repeat spacing from our original Cryo-TEM data.^9^ We then varied the depth of the interaction well to see whether we could further increase the stabilization of the stomatocyte with respect to the prolate spheroid (see the Supporting Information). One unforeseen consequence was that for very strong interactions, lamellar structures were formed instead of invaginations. Accordingly, we chose a value where the previously observed geometry remained stable within our simulation time. The results are shown in Figure 2 A, together with those of the initial model. They show that peptide-peptide interactions are, in principle, capable of stabilizing invaginations even against larger osmotic stress, requiring less excess membrane. However, the approach remains simplified. A more rigorous treatment of the screened electrostatic interaction between curved interfaces is very challenging to incorporate. Nevertheless, the present interaction potential already enhances the sorting to the negative mean and negative Gaussian curvature region of the invagination (see Supporting Information) and neck constriction, thereby bringing our modelling into closer agreement with the experimentally observed structures (stomatocyte examples shown in Figure S11, B).

**Figure 2:**
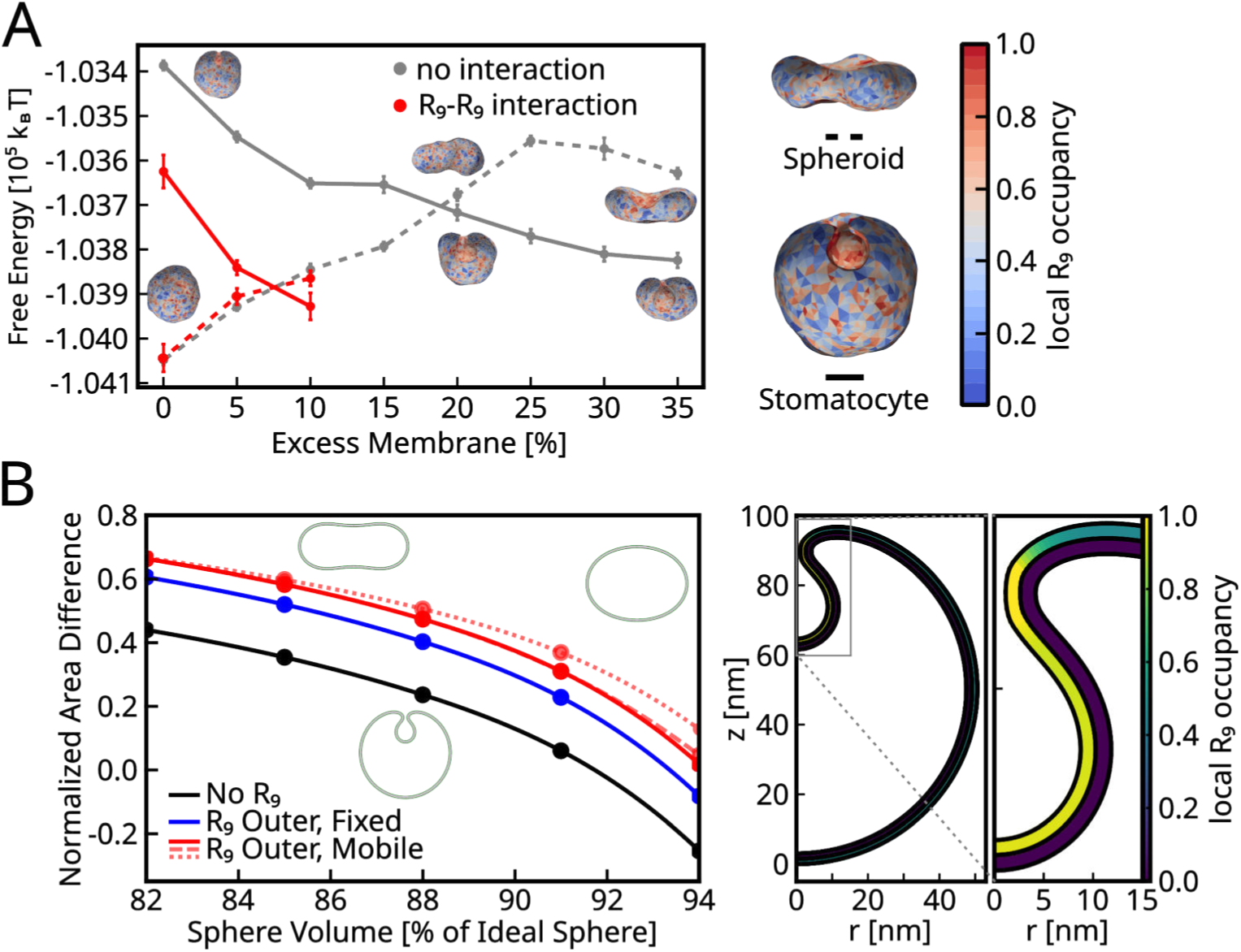
**(A) Monte Carlo Simulations Left**: Relative stability between spheroidal and stomatocyte-like morphologies in the presence (red) and absence (grey) of the interaction potential (*ɛ* = 0.0112 kT*/*nm^2^). Simulation snapshots illustrate representative shapes at corresponding reduced volumes. Error bars denote the standard error of the mean (SEM) with 95% confidence intervals. Right: Representative picture of a spheroid and stomatocyte-like structure with color map indicating the average R_9_ occupancy. (B) **Continuum Model Left**: Phase diagram for single-component systems in the (*ν, m*_0_) parameter space, following the inbudding transition. The coexistence curve (lines), interpolated with a cubic spline, separates the oblate and inbudded states from above and below, respectively, and is derived from sampled data points (circles). The diagram reflects variations in conditions such as the presence of R_9_ on the outer leaflet (R_9_ Outer; blue), its redistribution across the membrane (R_9_ Outer, Mobile; solid red line), and neck adhesion, influenced by different binding energies *ɛ* due to R_9_ (R_9_ Outer, Mobile, Sticky Neck; red dashed line for *ɛ* = 6 pN·nm^−1^ ; red dotted line for *ɛ* = 12 pN·nm^−1^). Right: Equilibrated inbudded structure for a single-component vesicle with a sticky neck (*ɛ* = 12 pN·nm^−1^) due to R_9_ -mediated membrane adhesion. The phase parameters are *ν* = 0.91 and *m*_0_*/*4*π* = 0.3. R_9_ is distributed over the outer leaflet. The inbud structure is emphasized in a close-up view with heatmap indicating local R_9_ occupancy.

#### II. Continuum Model

Building on the extended Helfrich–Hamm–Kozlov formalism^40,42,43^ and the continuum model of Ryham et al.,^44–46^ we developed an elastic framework for enclosed vesicles under physiologically relevant constraints.^47^ The model incorporates finite bilayer thickness, incompressibility, and constant inner water volume, consistent with experimental conditions.^48,49^ The equilibrium morphology of a 50 nm LUV is determined by the leaflet area asymmetry and the relative water content, with phase boundaries obtained by comparing the energies of the ”inbudded” and ”unbudded” states (see SI for further details). To disentangle the contributions of different physical mechanisms, we considered a hierarchy of models with increasing biophysical detail. Single-component vesicles, representing membranes without lipid redistribution, were simulated with the following conditions (see Fig. 2 B, left): no R_9_, immobile R_9_, mobile R_9_ on the outer leaflet, and mobile R_9_ with additional neck adhesion. Multi-component vesicles, which allow lipid mixing, were modeled under analogous conditions (see Fig. S8).

The results show that lipid mixing markedly enhances the tendency for inbudding, reflected by an upward shift of the coexistence line. This effect can be interpreted as a renormalization of the physical parameters; as in the the area-difference elasticity (ADE) model, demixing effectively softens the membrane while simultaneously adjusting the spontaneous curvature and asymmetry to stabilize the inbuds.^47,50,51^ Among R_9_-related mechanisms, induction of negative spontaneous curvature is the dominant factor, followed by the mobility of R_9_, which enables preferential accumulation at the inbud (reaching ∼ 80-90% occupancy compared to 50% in the leaflet bulk, see Fig. 2 B, right). Finally, neck adhesion contributes only in the slim-neck regime, which occurs at high relative water content when the inbud and its neck are small; in this regime, adhesion helps stabilize narrow necks, but does not drive their formation.

### Arginine-rich Peptides Reorganize Lipids Leading to Membrane Remodelling

The simulation results presented in previous sections point to a clear effect of both the arginine peptide content and the membrane lipid composition on the peptide-membrane interactions. In particular, the presence of phosphatidylethanolamine (PE) and phosphatidylserine (PS) lipids drives the K_9_ and R_9_ membrane adsorption and the peptide insertion within the membrane, resulting in local lipid density changes within the membrane. Here, we experimentally validate both findings by combining analysis of Laurdan fluorescence spectra and cryo-TEM imaging of large unilamellar vesicles (LUVs) of the three lipid compositions interacting with either K_9_, R_4_, or R_9_.

Laurdan, a membrane sensitive fluorescent dye, is usually utilized to monitor changes in the membrane phospholipid phase transitions,^52,53^ due to the sensitivity of its emission spectra to the membrane lipid packing (which arises from the non-lipid local environment of the naphthalene fluorescence moiety, residing near the glycerol backbone of a lipid bi- layer.^21^) Therefore, Laurdan emission spectra can vary due to peptide interactions with lipid headgroups, as well as due to peptide locally modifying lipid packing. Indeed, we observe that the Laurdan general polarization (GP) changes upon addition of peptides to liposomes (Fig. 3, top panel). As a starter, our results show almost no peptide effect on the membrane of POPC-containing LUVs. The impact of tetraarginine R_4_ on the negatively charged LUVs is similarly negligible. However, nonalysine and nonaarginine change the Laurdan GP of negatively charged bilayers significantly, pointing to a substantial perturbation of the interfacial/glycerol region, consistent with partial insertion and/or lipid reorganization (as suggested by simulation results, see Figure S1 and S2). Nonaarginine interacts more strongly with negatively charged membranes than nonalysine, as implied by the larger ΔGP change induced, in agreement with our MD results.

**Figure 3:**
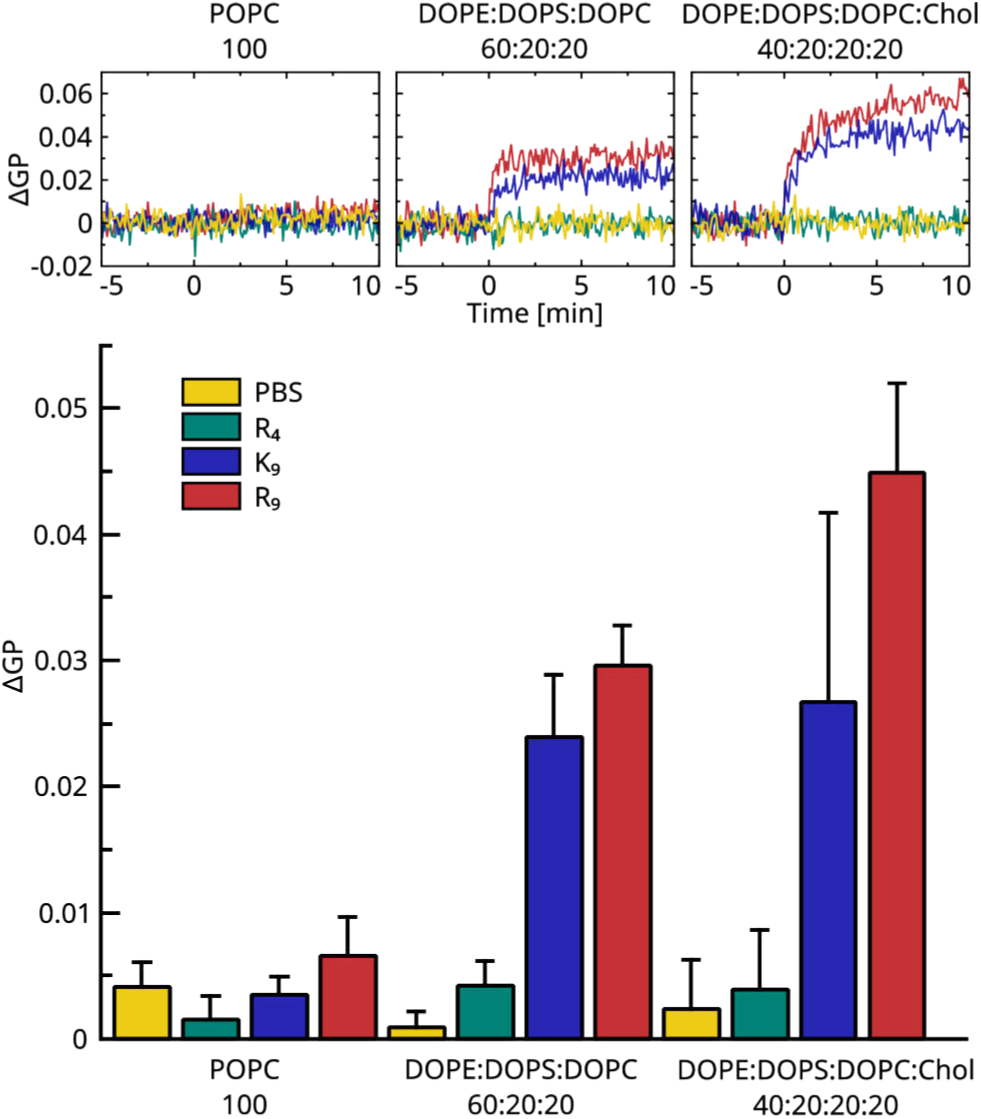
Peptide effect on various lipid membranes depicted as changes of a general polarisation factor (ΔGP) of laurdan spectra after adding R_4_ (green), K_9_ (blue) or R_9_ (red) peptides or no peptide (PBS -/- buffer control; yellow) to the LUV suspension at time 0 min. Top: Examples of ΔGP time evolution for three different LUV membrane compositions: POPC (left), DDD (middle), and DDDC (right). Bottom: The average GP increase for three independent measurements after peptide addition.

Similar results were obtained for the system prepared in PBS +/+ buffer containing Ca^2+^ and Mg^2+^ ions, approaching biologically more relevant conditions (see Fig. S4). Furthermore, Laurdan can be incorporated into the membrane of extracellular vesicles (EVs) extracted from ovarian cancer cells OVCAR-3, which are similar in size to LUVs. Figure S5 shows that, in accord with LUV results, R_4_ induces only minor GP changes, and the effect of nonalysine on the EV membrane is smaller than that of nonaarginine.

For all three investigated lipid compositions (exactly mirroring those employed in MD simulations), cryo-ET allows us to observe that LUVs without the presence of peptides, incubated in PBS (-/-), display a characteristic spherical shape, with occasional multivesicular LUVs present in the dispersion. In contrast, incubation with peptides results in a complex landscape of LUV morphologies (Fig. 4 A), broadly categorized as either the membrane remodelling, occurring at the membrane of a single LUV, or the intervesicular interaction, resulting from peptide-induced aggregation of multiple vesicles.

**Figure 4:**
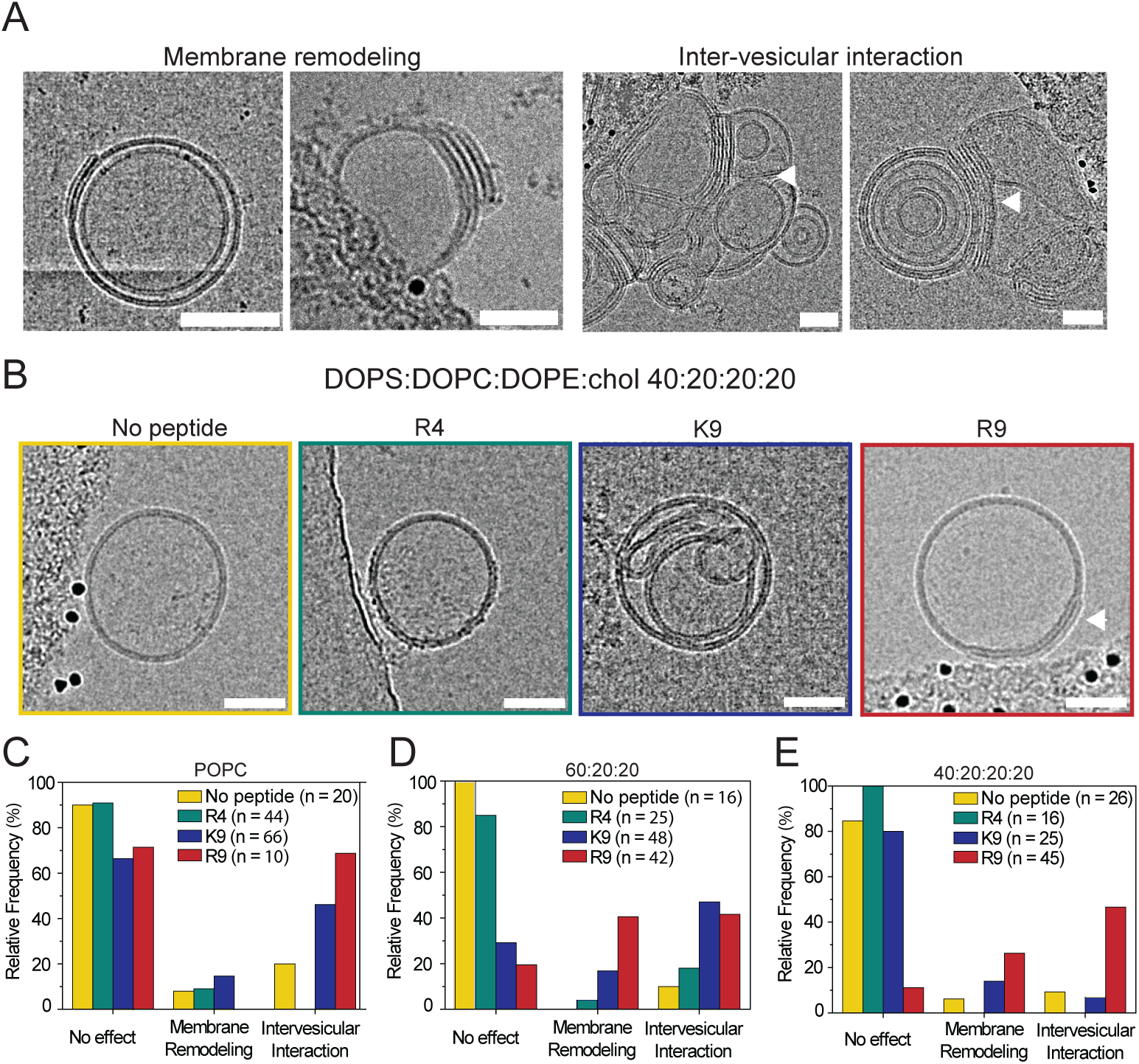
Peptide-membrane interaction shows a complex landscape of morphologies and intervesicular affinity modulated by peptide type and lipid composition. (A) Examples of cryo-EM images of the observed morphologies of membrane remodelling (changes in the lipid bilayer shape or contour upon interaction with peptide) and intervesicular interaction of LUVs upon incubation with peptides. (B) Representative cryo-TEM images of DDDC LUVs interacting with R_9_ (red), K_9_ (blue), R_4_ (green), or no peptide (PBS -/-, yellow). (C-E) Occurrence of observed vesicular morphologies for either R_9_ (red), K_9_ (blue), R_4_ (green), or no peptide (PBS -/-, yellow) interacting with POPC LUVs (C), DDD LUVs (D), or DDDC LUVs (E). The various morphologies or features can appear simultaneously on the same vesicle. Scale bar, 50 nm.

We observe that the occurrence and intensity of such morphologies are highly dependent on the peptide and LUV lipid composition (Fig. 4 B-E and Fig. S9), in line with the Laurdan measurements, MD simulations, and previous reports.^9^ In the case of cholesterol-containing systems (Fig. 4, B), we observe that R_9_ drives significant aggregation and membrane remodelling, including previously reported multilamellarity (Fig. 4, B), with only a small fraction of vesicles showing no detectable effect. In contrast, incubation with K_9_ or R_4_ results mainly in intervesicular interaction, with very limited membrane remodelling. LUVs composed exclusively of POPC show a markedly different landscape (Fig. 4, C). The peptide presence induces exclusively aggregation and only minimal membrane remodelling. For PS and PE-rich systems not containing cholesterol (DDD), we observe a similar trend as for the DDDC LUVs (Fig. 4, D), but with a more pronounced trend towards intervesicular interaction (aggregation and adhesion).

Overall, the cryo-TEM analysis reveals that the lipid composition is indeed essential for the arginine-rich peptides’ mechanism of action, and that the specific effect of R_9_ on membranes is an extensive remodelling of PS and PE-rich membranes leading to membrane folding, while other cationic peptides result in nonspecific aggregation. While the presence of PS and PE is necessary for the rearrangement of membranes, the presence of cholesterol appears to slightly reduce the extent of such remodelling.

### Cryo-EM Tomography on LUVs

Having identified the membrane lipid composition resulting in the most efficient peptide interaction, we then performed cryo-ET on the cholesterol-containing DDDC liposomes incubated with R_9_ at different time points (namely 1, 3, 7, and 10 minutes) to visualize the membrane remodelling process at higher resolution. The obtained images were then compared to those of control LUVs (LUVs without peptide presence, see Fig. 5 A, B) to determine i) how specific morphologies appear over time and ii) what the specific action of the arginine-rich peptides is.

**Figure 5:**
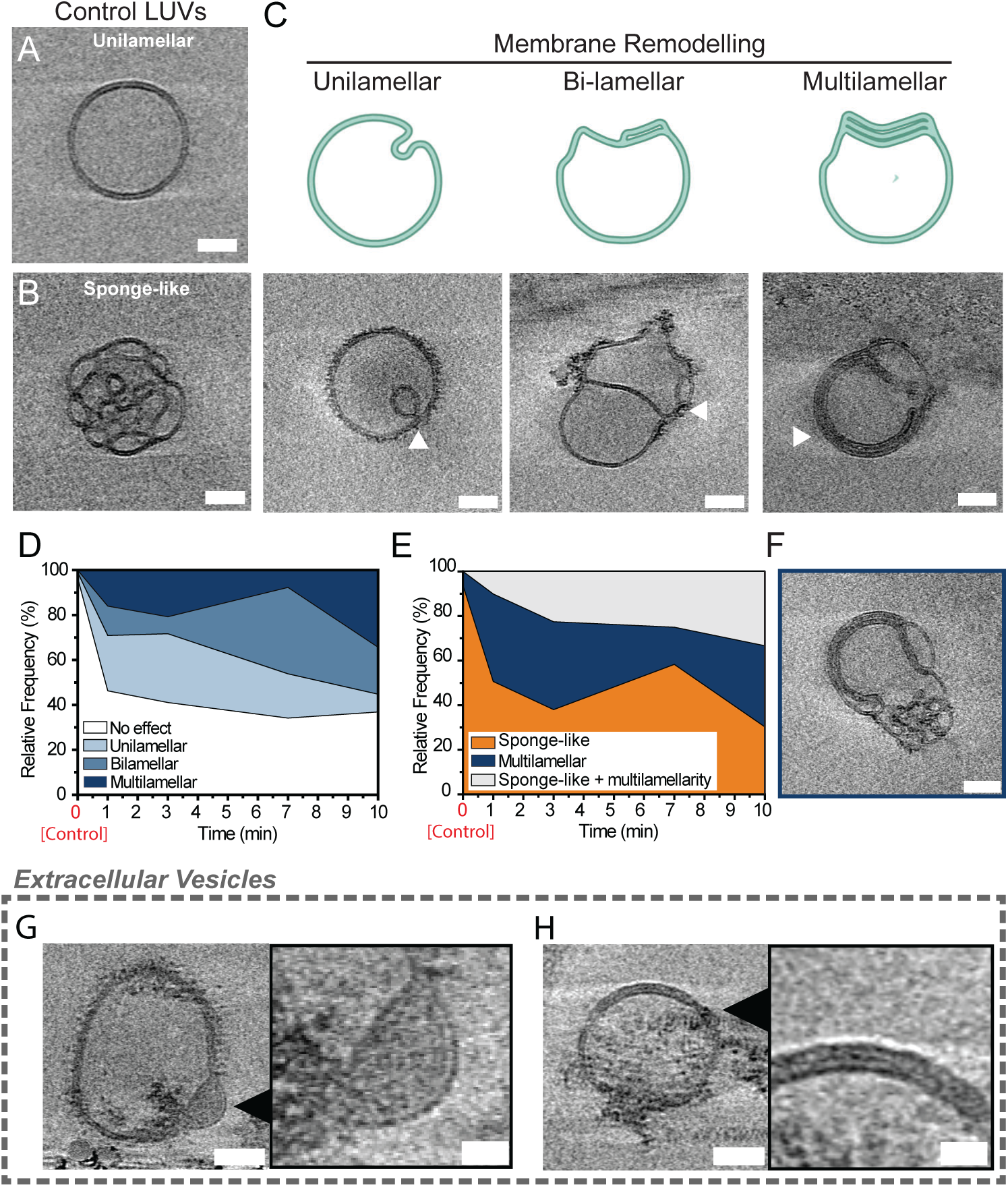
R_9_ remodels membranes by favoring membrane folding and stacking. (A) Representative tomographic slice of cholesterol-containing DDDC LUVs displaying the characteristic spherical and unilamellar morphology. (B) Representative tomographic slice of control LUVs displaying a sponge-like inner structure, probably occurring due to high PE content. (C) Summary of the membrane-remodelling morphology classification, including unilamellar membrane remodelling, such as budding, bilamellar remodelling, which includes bifurcation or droplets, and multilamellar remodelling, which occurs only when three or more bilayers are stacked. The yellow triangles in representative cryo-ET images indicate the specific morphology. (D) Quantification of remodelling class occurrence at different incubation time points obtained from cryo-ET images. (E) Quantification of sponge-like and multilamellarity co-occurrence at different incubation times as extrapolated from cryo-ET tomographies. (F) Representative cryo-ET slice of liposomes displaying the co-existence of a sponge-like structure and multilamellarity. For all images scale bar is 50 nm. (G-H) Representative cryo-ET images of EVs incubated with R_9_ for 10 min resulting in clear membrane bilamellarity (blue triangles) with zoomed-in detail on the remodelling spot (blue squares, scale bar: 20 nm). In most cases, the two bilayers are difficult to distinguish due to the lower contrast caused by the presence of proteins and glycans on the membrane.

Extending the imaging to the 3D space revealed several important, otherwise undetectable, aspects. In general, we classify the various types of vesicle morphology caused by peptide-membrane interaction depending on the number of bilayers involved (Fig. 5 C). Unilamellar interactions, such as inward budding, result in a folded membrane but without the formation of stacked bilayers. Bilamellar morphologies are characterized by membrane bifurcations, while the multilamellarity is here defined as three or more bilayers stacked together.

In the case of control LUVs (without peptide), we observe two main morphologies: a majority of (expected) spherical unilamellar vesicles (∼ 85% of the LUV population, Fig. 5 A), and a smaller fraction (15%) displaying an inner membrane network, similar to water channels of continuous cubic phases or inverse hexagonal phase (Fig. 5 B). This feature is most likely due to the high PE content within individual vesicles, as PE favors highly negative curvatures; so the membranes rearrange to accommodate the lipid-specific geometry.

In contrast, interaction with R_9_ results in significant remodelling across all three typologies (Fig. 5 C), with multilamellar features being predominant (∼ 34% at a 10 min incubation time). Analysis of the occurrences of each morphology over time reveals that multilamellarity increases over time at the expense of both uni- and bi-lamellarity (Fig. 5 D). Similarly to untreated LUVs, those interacting with R_9_ still display a sub-population containing an inner membrane network. Interestingly, further analysis on the co-occurrence of such structures with the remodelling morphologies shows that multilamellarity occurs more often together with the inner network, and that such co-existence increases over time until 7 min of incubation and declines afterwards, while the number of multilamellar-only vesicles increases (Fig. 5 E, F). The time evolution of the different morphologies suggests that R_9_ remodels membranes with dramatic membrane bending and folding, resulting in either inward budding or multiple membrane stacks. Specifically, the peptide action favors stacked morphologies and, in some instances, highly stacked bilayers. Examples of increased multilamellarity in time, possibly utilising the excess inner network membrane, are shown in Figure S11, A, while Figure S11, B shows observed nonaarginine-induced inward budding structures.

Increasing the membrane’s biological complexity by incubating the peptide with EVs, rather than LUVs, leads to similar remodelling morphologies (Fig. 5 G, H, and Fig. S10), with examples of membrane bilamellarity comparable to those occuring on LUVs. The membrane bilamellarity on EVs can occur both as distant bilayers (Fig. 5 G) and stacked membranes (Fig. 5 H). In addition, cases of no visible effect of the peptide (Fig. S10 B) as well as ”burst” vesicles (Fig. S10 C) are observed, too.

### Cryo-ET Analysis by Machine Learning

Reconstructed tomograms were cropped to contain LUV vesicles, cut into *z*-slices, and each slice was processed by a two-step convolutional neural network classification procedure for properties such as R_9_ presence on the vesicle membrane or membrane multilamellarity (see Fig. S12). The results for slices belonging to individual vesicles were then evaluated together. Finally, each lipid vesicle was assigned a percentage reflecting how much of the property of interest the given vesicle manifests. For example, 0% multilamellarity means that the given vesicle is strictly unilamellar, while 80% multilamellarity suggests that 80% of the membrane detected in this vesicle is multilamellar.

The amount of R_9_ peptide on the outer vesicle membrane for individual incubation time ensembles is shown in Figure 6, A, left, with examples of machine learning clasifications on the right (no R_9_ on the membrane in blue, nonaarginine on the outer vesicle membrane in red). Already with as short as 1 min R_9_ incubation with LUVs, nonaarginine is observed to reside on the membrane. Its membrane coverage increases with time, as shown in the 3 min and 7 min incubation datasets. The 10 min dataset shows a significant decrease in nonaarginine on the vesicle membrane. This can be caused by R_9_ inducing multilamellarity, penetrating inside vesicles (see Figure S11), or even causing vesicles to burst. The small nonzero value of R_9_ presence on the LUV membrane in the systems without R_9_ incubation can be considered as the error margin of this analysis.

**Figure 6:**
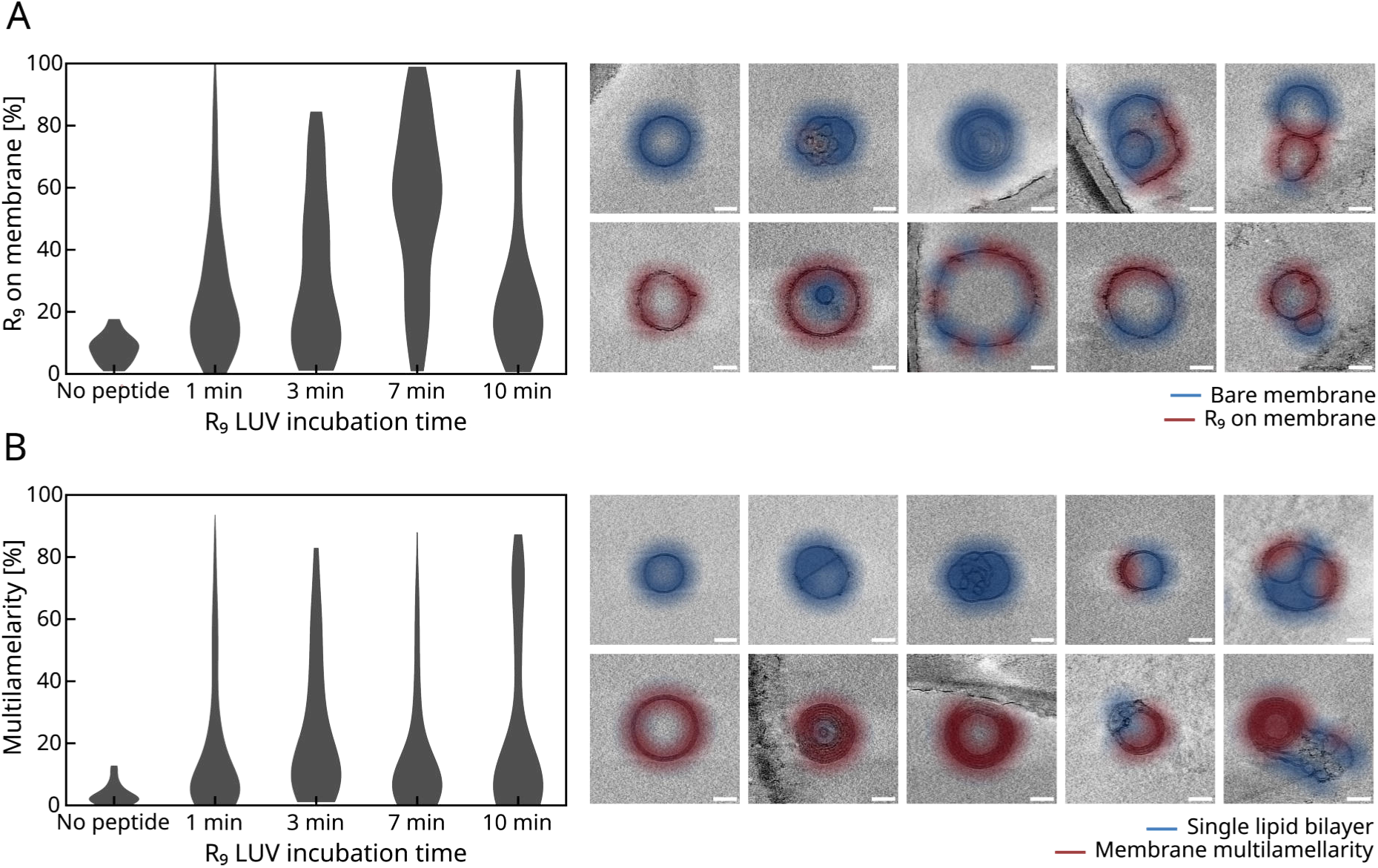
Cryo-ET LUV data processed by machine learning. (A) Left: The amount of R_9_ on the vesicle outside membrane detected in DDDC LUV vesicles without R_9_, as well as at various R_9_ incubation durations. Right: Examples of machine learning algorithm detecting nonaarginine on the membrane (red) as a contrast to the bare membrane (blue). Note that several species exhibit both. (B) Similarly, time evolution for multilamellarity occuence (left), with examples (right). Here, unilamellar membrane parts are shown in blue and multilamellar red. The scalebar is 50 nm.

**Figure 7:**
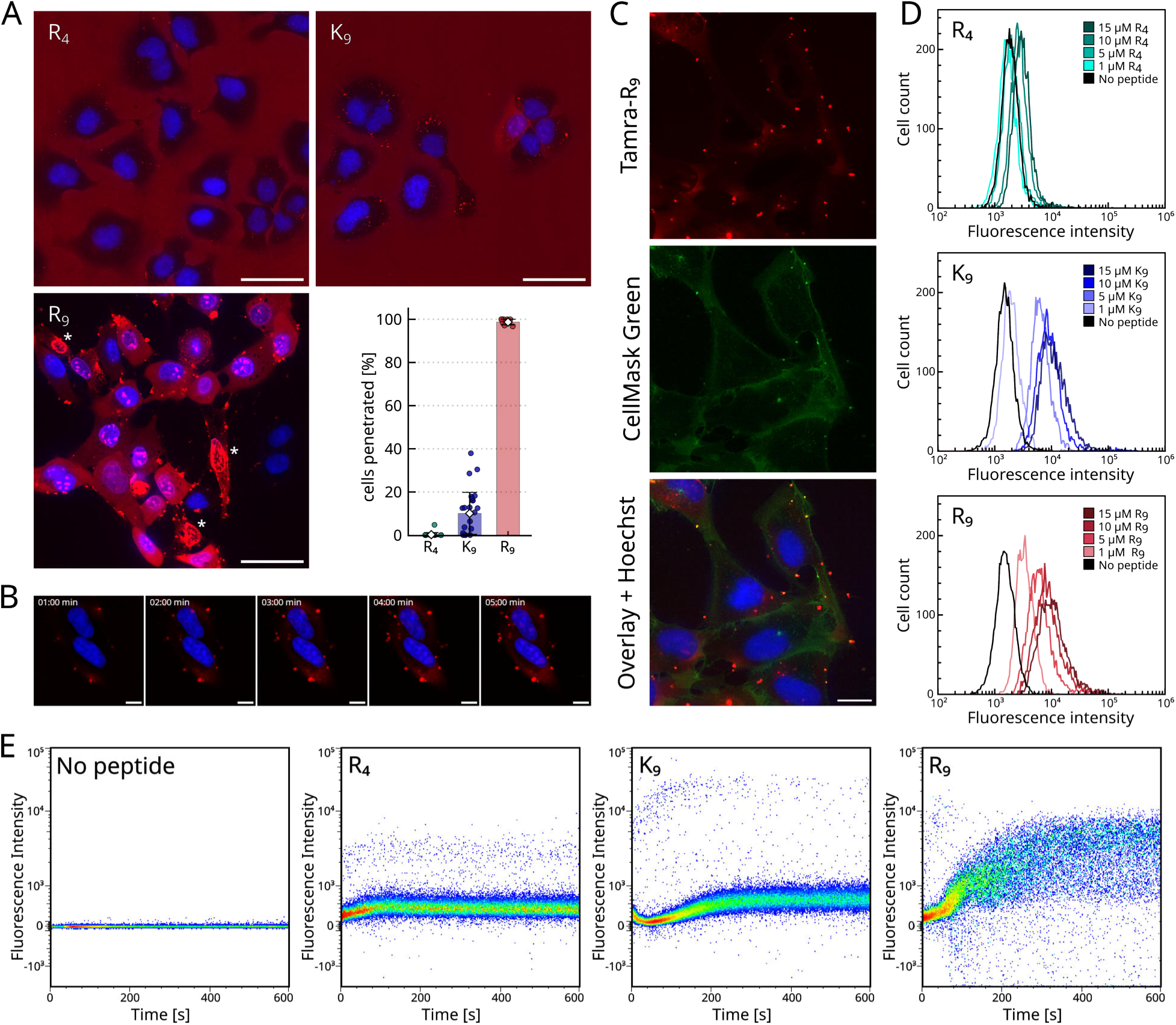
Nonaarginine translocates into U-2 OS cells. (A) U-2 OS cells incubated for 20 min with 10 *µ*M peptides: R_4_ (top left), K_9_ (top right) and R_9_ (bottom left), with white asterisks labelling dead cells. Tamra-labelled peptides in red, Hoechst-marked cell nuclei in blue. The scalebar is 50 *µ*m. Bottom right - a weighted average of penetrated cells from several technical and biological repeats. (B) Example of R_9_ cell-translocation timelapse. Tamra-labelled 10 *µ*M nonaarginine (red) was added to the cells at time 0 min. Cell nuclei labelled by Hoechst (blue). The scalebar is 10 *µ*m. (C) Nonaarginine and membrane collocalisation in the puncta on cell surface. 7.5 *µ*M Tamra-R_9_ is shown in red (top) and CellMask Green is visualising cell membranes (middle). Merged channels, along with Hoechst-labelled nuclei in blue showing the collocalization of R_9_ and membrane in the puncta (bottom). The scalebar is 20 *µ*m. (D) Flow cytometry Tamra-peptide fluorescence intensity histograms after 20 min peptide incubation. Cells without peptide (in black) and peptide concentrations of 1 *µ*M, 5 *µ*M, 10 *µ*M and 15 *µ*M were used. R_4_ (top left), K_9_ (top right) and R_9_ (bottom left). (E) Time-resolved flow cytometry with 1 *µ*M peptide added at 0 min. Fluorescence intensity time evolution with no peptide present with cells (left), R_4_ (middle left), K_9_ (middle right), and R_9_ is shown in right.

Figure 6, B, left, illustrates the time evolution of vesicle membrane multilamellarity, measured in datasets with varying time of R_9_ LUV incubation. Unilamellarity (blue) versus multilamellarity (red) neural network detection examples are shown in Figure 6, B, right.

Here, only a slight membrane multilamellarity increase occurs over time. Since the sponge-like vesicle population (see Figure 5, B) is a minority (15%), there is not that much free membrane to form the multilamellar structures.

### Nonaarginines Translocating into Cells

#### Fluorescence Microscopy

Fig. 7, A shows a comparison of cell penetration efficiency between tetraarginine, nonalysine and nonaarginine. U-2 OS cells were incubated with 10 *µ*M peptide for 20 min. This concentration was chosen such as to induce the passive cell translocation by R_9_, but not stress cells to the point of rapid death (see Fig. S13). Illustrative snapshots for R_4_, K_9_, and R_9_ are shown on top left, top right, and bottom left, respectively. Here, the highest contrast objects are dead cells, labelled by a white asterisk. In the bottom right, the calculated weighted average of penetrated cells shows almost no cell penetration in the case of R_4_ (0.2 ± 1.0%), while 10.1 ± 9.9% of the cells were penetrated by K_9_, and 98.7 ± 1.4% by R_9_. Shortly after R_9_ peptide addition to the U-2 OS cells (∼ 1 min), distinct peptide aggregates appear on the cell membrane (see Fig. 7, B). Similar shapes were also observed in HELA^8,54,55^ and Jurkat cells^8^ after the polyarginine addition. These R_9_-induced puncta are actually peptide-membrane complexes, as proven by collocalization of the Tamra-R_9_ channel and membrane Cellmask Green stain (Fig. 7, C) and further studied by CLEM in this article. Supplementary video S1 shows that these peptide-membrane puncta become a place of peptide entrance to the cell, with a visible influx of Tamra-labelled peptide into the cell (estimated peptide addition set to 0 min). Mere minutes after R_9_ enters the cell, it transports into the nucleus and stains DNA (see Fig. 7, B, Supplementary video S1), as was also observed for the HELA cell line.^56^

The more R_9_ is added, the more peptide-membrane puncta are created, so more tension is generated upon the cell membrane. Thus, at some point, the cells begin to slightly shrink. Since they do not have an infinite membrane supply buffer, if enough R_9_ is added (in our experience, around 15 *µ*M of Tamra-labelled R_9_ or more, see Fig. S13, right), a significant amount of cells rapidly burst and die. An example of burst cells is also visible in the Fig. 7, B, marked by white asterisks. Similarly, we observe these puncta in the case of K_9_, and much less in the case of R_4_. However, almost no cell penetration occurs in either case.

#### Flow Cytometry

Peptides exhibiting membrane association and translocation into cells produced increased cell-associated fluorescence together with a relative reduction in background contribution, see Figure 7, D, in accord with the microscopy observations in Figure 7, A. These effects yield well-resolved rightward shifts in fluorescence intensity distributions. Concentration-dependent fluorescence shifts provide a more robust metric of peptide–cell interaction than absolute fluorescence values. R_9_ and K_9_ induced pronounced concentration dependent fluorescence shifts, consistent with efficient cellular association. In contrast, a minimal change in fluorescence intensity occurs in the case of R_4_, indicating a weak interaction with U-2 OS cells.

Note the prominent drop in the number of selected cells-events (Fig. S14, A, first row) in the first R_9_ gate from 96% to 84% and their presence outside the selected gate, suggesting pronounced cell aggregation into multiplets (SSC-A and FCS-A) and changes in internal cell structure (SSC-A increase). A less prominent drop was also observed for K_9_ (to 92%). The cells and their aggregates outside the gated area were characteristic with very high PE-A values (*>* 3∗10^4^, Fig. S14, A, third row), demonstrating that the observed effect is associated with the high amount of peptide present and not the physiological state of the cells.

Since only single cells were gated, the contribution of possible aggregated multiplets is excluded from the fluorescence histograms (see Fig. S14, A). Similarly, cells that bursted and died from the exposure of high nonaarginine concentrations, as shown in Figure S13, right, do not contribute either. If included, the R_9_ fluorescence intensity histograms would shift even more to higher values.

Additionally, the fluorescent intensity histograms do not allow us to distinguish between fluorescently-labelled peptides interacting with the outer cell membranes and those translocated into cells. In fact, the translocated peptides’ signal might be slightly quenched compared to the extracellular environment of the outer membrane singnal, as quantified for Penetratin uptake.^57^ Therefore, the K_9_-membrane interaction, suggested by MD simulations (Fig. 1) and detected in LUV experiments (Fig. 3 and Fig S4, Fig. S5, Fig. 4), without significant cell translocation (observed in live-cell confocal imaging, Fig. 7), might explain similar nonaarginine and nonalysine histogram shifts in Fig. 7, D.

Time-resolved flow cytometry was performed using a much more sensitive experimental setup. First, peptide stability was checked (see Fig. S15). No big peptide aggregation or significant increase of fluorescence signal in time was observed in a system containing only peptides and HBSS buffer, in the absence of cells.

Immediately after 1 *µ*M of Tamra-labelled peptide was added to the U-2 OS cell suspension (see Figure 7, E), singlet cell population fluorescent signal increased. Cells incubated with tetraarginine show the lowest increase, suggesting low peptide-membrane interaction. Similarly, K_9_ also interacts weakly. Here, the temporary signal drop, occuring around 40 s, is probably an artefact caused by suboptimal sample mixing after peptide addition, showing local concentration inhomogeneity. In contrast to R_4_ and K_9_, the highest fluorescent signal is observed for singlet cell events incubated with R_9_, showing fast and excessive peptide adhesion to the singlet cell membrane.

The time-resolved flow cytometry gating and workflow is shown in Fig. S16. The aggregated events show similar fluorescence behaviour to singlet cell population, regardless of the peptide added. In contrast, the fluorescence intensity of subcellular-size particles is higher in all three cases, most prominently in the case of K_9_.

Additionally, note the absence of very small subcellular fragments for K_9_ and R_9_ (visible in Fig. S16, top row, bottom left part of the figures), in contrast to cells without peptides or R_4_. This can be explained by a peptide-induced fragment aggregation, previously observed by dynamic light scattering measurements on LUVs,^9^ showing R_9_-induced size increase in contrast to no change observed with R_4_.

#### Cells Observed by CLEM

Following the imaging of peptides entering cells, we aimed to directly visualize in situ the interaction of cells with R_9_ in order to understand how nonaarginines reshape cells, thus potentially decipher their entry mechanism. To do this, we employed a correlative approach to live-cell imaging using the Tamra-labelled R_9_. We first prepared cryo-EM-suitable samples of cells incubated with Tamra-R_9_ by plunge-freezing in liquid ethane. Subsequently, the samples were visualized via cryo-fluorescence microscopy (see Fig. 8 A, B), confirming the presence of fluorescence peptide puncta, located within and on the periphery of the cell (Fig. 8 B), which corresponded to features visible in cryoEM (Fig. 8 C). Having identified Tamra-R_9_ puncta of interest, we performed cryoET acquisition at the exact location by correlating the fluorescence and EM images. Using this approach, we were able to obtain a three-dimensional view of the membrane remodelling at the cell surface containing peptide clusters before entry.

**Figure 8:**
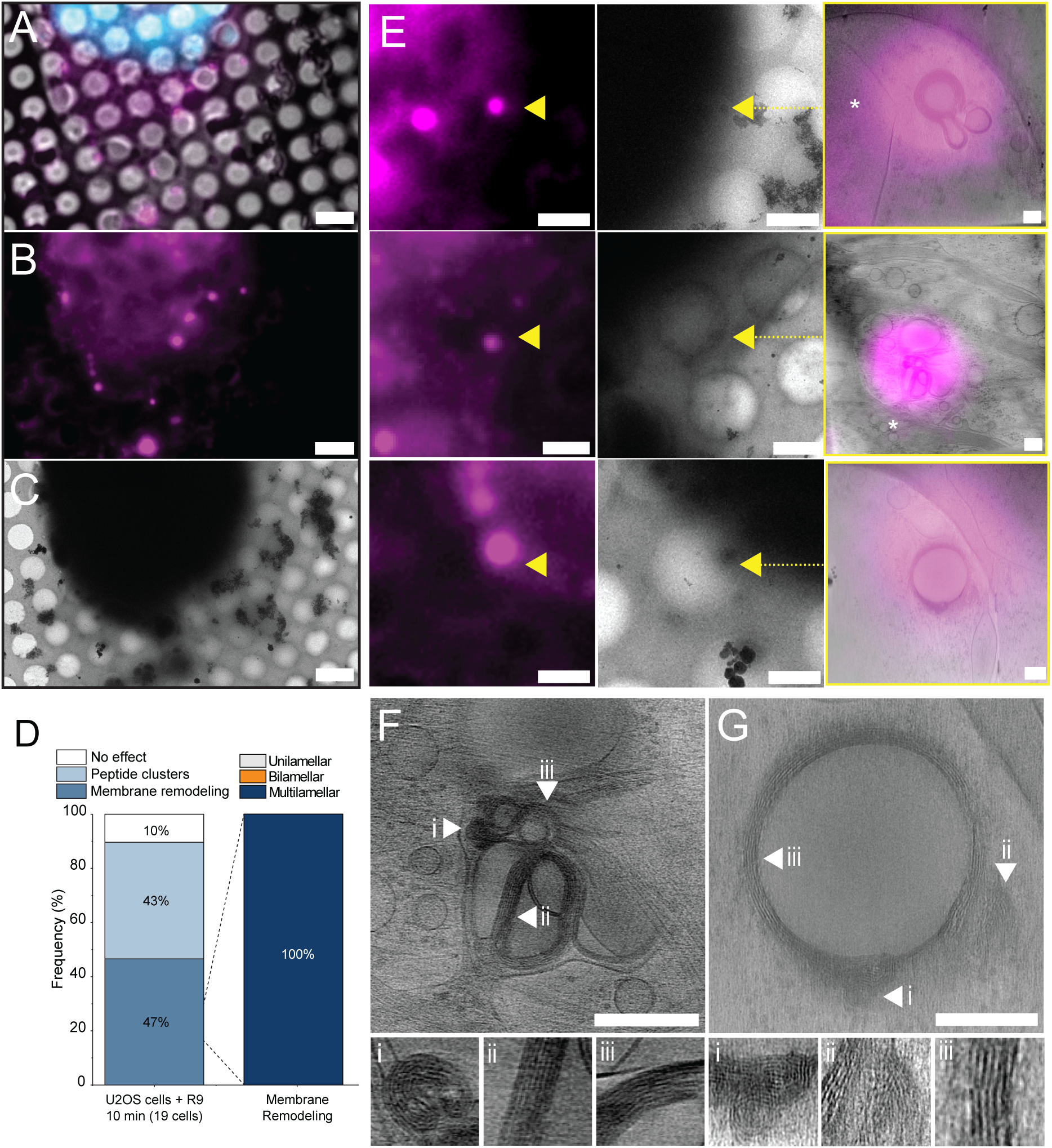
R_9_ interacting with cell membranes results in extensive multilamellarity. (A-B) Representative cryoFM image of U-2 OS cell growing on an EM grid incubated with Tamra-labelled R_9_, presented both as overlay of brightfield and fluorescence signal (A) and peptide-only signal (B). Magenta represents Tamra fluorescence, and cyan is the cell nucleus stained with Hoechst before plunge-freezing. (C) Cryo-electron microscopy image of the same sample coordinates presented in A and B. Scale bar: 5 *µ*m. (D) Statistical analysis of type of morphology observed across peptide-positive puncta. (E) Representative correlations of peptide puncta, displaying membrane remodelling and multilamellarity, presented as peptide-only fluorescence (left column), corresponding electron microscopy image (middle column), and cryo tomography image with overlayed peptide fluorescence signal (right column). Scale bar: 2 *µ*m. (F-G) Representative tomographic slices displaying R_9_-induced membrane remodelling, with examples highlighted by white triangles and corresponding insets. Scale bar 200 nm.

We observe two primary morphologies across the R_9_-positive puncta, namely membrane remodelling and peptide clustering (see Fig. S17), which occur at 47% and 43% of the tomographs, respectively (Fig. 8, D, as obtained from 18 different cells from a single CLEM experiment). Within the membrane remodelling morphology, all the visualized puncta present clear and extensive multilamellarity and membrane folding (Fig. 8, D), with distinct examples where we were able to observe the multilamellar feature directly corresponding to the fluorescence signal and connected to the cell plasma membrane (Fig. 8, E). This membrane remodelling occurs in a form of membrane vesicle structures, either directly pinching off the cell membrane (Fig. 8, F), or proximal to the plasma (Fig. 8, G), displaying a large amount of stacked bilayers (Fig. 8 F, G insets). In addition, we also observe multilamellarity in between multiple vesicles (probably arising from the adhesion of the cells on the EM grid mesh), which are strongly aggregating and displaying multilamellar morphologies at the vesicle-vesicle interaction sites (Fig. S18).

Interestingly, the effects of R_9_ interaction in cells appear very prominent and clearly visible despite the peptide concentration being ∼ 10 times lower than that for both LUVs and EVs (10 *µ*M for cells as opposed to ∼ 150 *µ*M for LUVs and EVs cryo-EM). Note that for cell-peptide experiments, the upper limit for R_9_ concentration is dictated by having a wide enough time window before peptide-induced cell death (see Figure S13, right). Nevertheless, the observed R_9_ effect on the cell membrane is morphologically similar to what we observed for R_9_-LUV interactions (i.e, vesicular aggregation, multilamellarity, and membrane folding), although we do not observe topologies with only unilamellar or bilamellar remodelling.

The presence of only multilamellar events could be both due to the small dataset probed (27 acquired tomograms within a single cryoCLEM experiment), specific interaction or biomolecular signature of the individual cell, or by detection limits of peptide fluorescence for less lamellar structure, since only cases of large enough amounts of membrane-bound R_9_ were selected. While we cannot exclude either scenario, statistical analysis of membrane remodelling occurence suggests that membrane remodelling is not specific to individual cells (Fig. S19). Moreover, considering the occurence of multilamellar events derived from LUVs-peptide cryoET (∼ 20% of all remodelling events), the probability of observing only membrane remodelling events by chance (under a binomial model) is approximately 1.05x10^−1^^9^ (*p <* 10^−1^^8^, one-sided binomial test). This strongly suggests that the membrane remodelling induced by R_9_ on cells is specifically multilamellar, rather than a random mixture of sub-morphologies.

## Conclusions

We employed a battery of techniques ranging from fluorescence microscopy and cryoelectron tomography to atomistic molecular dynamics simulations and continuum modelling to unravel the modes of action of oligoarginines as iconic cell-penetrating peptides in passively penetrating phospholipid membranes.^1,58^ To this end, we investigated systems of increasing compositional and structural complexity, from large unilamellar vesicles over extracellular vesicles to live cells.

First, by comparing the action of nonaarginine to that of tetraarginine and nonalysine, we confirm that both length and chemical composition of the cationic peptide matter, with only the former species being an effective cell-penetrating peptide. Second, simulations show that as a first step of penetration, nonaarginine accumulates at the membrane leading to lipid reorganization and bilayer deformations, particularly when negatively charged (PS) and curvature-inducing (PE) lipids are present. This is consistent with our fluorescence microscopy observations of the peptide-membrane aggregates (puncta) preceding cell penetration, as well as the deformed bilayer structures, including budding, bifurcations, and multilamellarity, revealed by cryoelectron tomography. Third, our cryo-ET imaging, aided by machine learning analysis, reveals that the morphological landscape of membrane remodelling narrows down as the complexity of the membrane models increases. In LUVs, which are composed of only three phospholipids and cholesterol, membrane reorganization involves both unilamellar deformations, such as budding, and formation of double or multiple membrane layers. In biological membranes, namely EVs and cells, the landscape decreases to simpler types of remodelling, either bilamellar (EVs) or highly multilamellar (cells), due to the interaction with R_9_.

Taken together, our results indicate a single mode of action of R_9_ on the membrane, which is then modulated by the complexity and accessibility of the membrane. R_9_ preferentially binds to membranes containing negatively charged lipids, with subsequent reorganization of lipids around peptides resulting in a local membrane curvature, as demonstrated by MD simulations and theoretical modelling.

Such local spontaneous curvature then drives unilamellar remodelling (like the membrane budding observed in both theoretical modelling and cryo-ET), and, subsequently, bilamellarity or multilamellarity is formed due to peptide-mediated adhesion between membranes. This ”fold and stack” mechanism is the primary mode of interaction of R_9_ with membranes (see Fig. 9). When the membrane reservoir is limited, as in the case of single LUVs and EVs, R_9_ action can only result in up to bilamellar structures with individual vesicles alone. However, if more membrane becomes accessible, as in the case of inter-vesicular interaction or additional reservoir of a sponge-phase, multilamellarity can arise. This is precisely the source of the broad morphological landscape observed in LUVs. When there is sufficient unilamellar membrane to be folded, R_9_ addition results in highly multilamellar structures, as observed in cells via CLEM.

**Figure 9:**
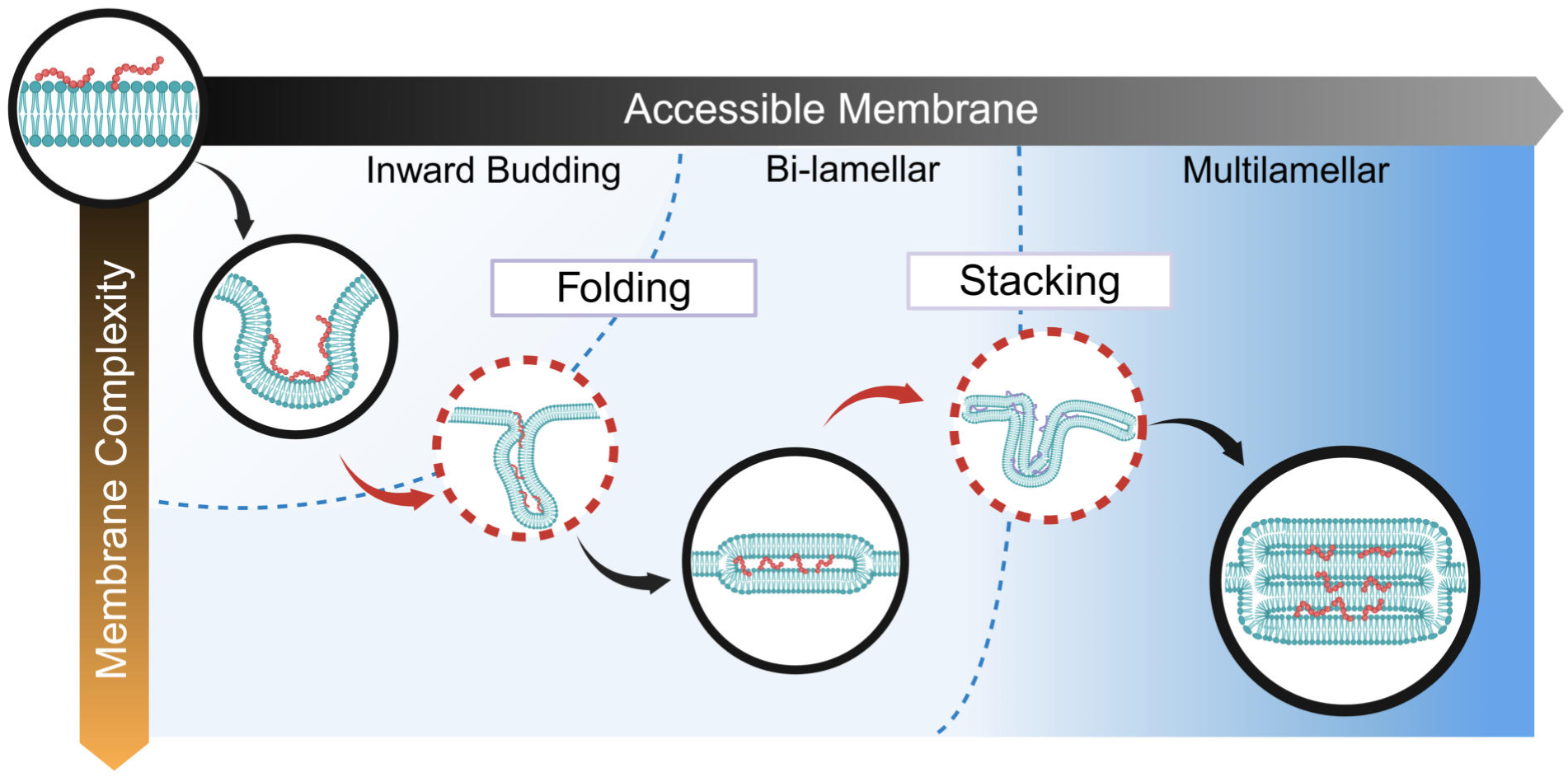
Proposed mechanism of R_9_-induced membrane remodelling leading to peptide cell translocation. Graphical illustration summarizing the proposed mode of action of R_9_ on membranes. Circles with solid black outlines represent stages experimentally observed in this work; circles with dashed red outlines are intermediate stages not experimentally observed and hypothesized based on indirect evidence. Peptide adsorption on the membrane induces lipid reorganization and spontaneous curvature, leading to initial budding. Such local curvature remodelling is then likely fused by the peptide, due to its high aggregation and fusogenic properties (”folding”), forming a bilamellar morphology. The process can then occur recursively on bilamellar membranes, unilamellar membranes, or both (”stacking”), yielding different degrees of multilamellarity. The amount of membrane reservoir the peptide can remodel is the limiting factor for achieving multilamellarity, whereas membrane composition and complexity enhance the peptide’s effect.

## Acknowledgement

PJ acknowledges support from the European Research Council via an ERC Advanced Grant no. 101095957. KB thanks the CB2 group for valuable discussions and support. She also acknowledges support from Charles University, where she is enrolled as a Ph.D. student, and HPCg at IOCB Prague for computational resources. M.V. acknowledges the Czech Science Foundation for support via grant number 25-16117S. M.I.M. acknowledges support from the EMBO Installation Grant no. 6030, and the Ministry of Education, Youth and Sports of the Czech Republic via an INTER-COST grant no. LUC25110.

O.A. also acknowledges funding from the Minerva Foundation with funding from the Federal German Ministry for Education and Research and the Minna James Heineman Foundation, the Henry Chanoch Krenter Institute for Biomedical Imaging and Genomics, the Schwartz Reisman Collaborative Science Program, and the Yeda-Sela Center for Basic Research. O.A. is an incumbent of the Miriam Berman presidential development chair. I.S. acknowledges support from Charles University, where he is enrolled as a Ph.D. student, Daniel Harries for valuable discussions, and HPCg at IOCB Prague for computational resources.

We acknowledge the access and services provided by the Imaging Centre at the European Molecular Biology Laboratory (EMBL IC), generously supported by the Boehringer Ingelheim Foundation. Financial support for access was provided by Instruct-ERIC (PID:25516) and iNEXT Discovery (PID:23416). We also acknowledge the CF CryoEM of CIISB, Instruct-CZ Centre, supported by MEYS CR (LM2023042) and European Regional Development Fund-Project UP CIISB No. CZ.02.1.01*/*0.0*/*0.0*/*18 046*/*0015974 through the CIISB funding (CIISB 220075C).

The authors thank A. Gálisová for providing a sample of HEK EVs for additional Laurdan experiment. The authors acknowledge the Imaging Methods Core Facility at BIOCEV, the institution supported by the MEYS CR (LM2023050 Czech-BioImaging), for their support and assistance with flow cytometry.

## Supporting Information Available

Further details on the computational and experimental procedures and results.

## Notes

### Competing Interest Statement

The authors have declared no competing interest.

