## Supplementary material for "Direct Membrane Penetration of Oligoarginines by Fluorescence and Cryo-electron Microscopy Combined with Molecular Simulations": SI

‡*Present address: Biochemistry Center (BZH), Heidelberg University, Heidelberg, Germany*

### Umbrella Sampling Simulations

#### Force Constants

Table S1: **Umbrella windows force constants** [kJmol$^{-1}$nm$^{-2}$] - for each umbrella window marked by membrane-peptide center of mass z-distance $d$ defining minimum of a harmonic potential.

| | POPC | | | DDD | | | DDDC | | |
|---|---|---|---|---|---|---|---|---|---|
| $d$ [Å] | $R_4$ | $R_9$ | $K_9$ | $R_4$ | $R_9$ | $K_9$ | $R_4$ | $R_9$ | $K_9$ |
| 1.45 | 1000 | 1000 | 1000 | 1000 | 1000 | 1000 | 1000 | 1000 | 1000 |
| 1.60 | 1000 | 1000 | 1000 | 1000 | 1000 | 1000 | 1000 | 1000 | 1000 |
| 1.75 | 500 | 1000 | 1000 | 1000 | 1000 | 1000 | 1000 | 1000 | 1000 |
| 1.90 | 500 | 1000 | 1000 | 500 | 1000 | 1000 | 1000 | 1000 | 1000 |
| 2.05 | 500 | 1000 | 1000 | 500 | 1000 | 1000 | 1000 | 1000 | 1000 |
| 2.20 | 500 | 500 | 1000 | 500 | 500 | 1000 | 500 | 1000 | 1000 |
| 2.35 | 500 | 1000 | 1000 | 500 | 500 | 1000 | 500 | 1000 | 1000 |
| 2.50 | 500 | 500 | 500 | 500 | 1000 | 1000 | 500 | 500 | 1000 |
| 2.65 | 500 | 500 | 1000 | 1000 | 1000 | 1000 | 500 | 500 | 1000 |
| 2.80 | 500 | 250 | 250 | 1000 | 1000 | 500 | 500 | 500 | 500 |
| 2.95 | 500 | 250 | 250 | 1000 | 1000 | 500 | 1000 | 500 | 500 |
| 3.10 | 500 | 250 | 250 | 500 | 500 | 500 | 500 | 500 | 500 |
| 3.25 | 250 | 250 | 250 | 500 | 500 | 500 | 500 | 500 | 500 |
| 3.40 | 250 | 250 | 250 | 250 | 500 | 500 | 250 | 500 | 500 |
| 3.55 | 250 | 250 | 100 | 250 | 500 | 500 | 250 | 500 | 500 |
| 3.70 | 250 | 100 | 100 | 100 | 250 | 250 | 250 | 500 | 250 |
| 3.85 | 100 | 100 | 100 | 100 | 250 | 250 | 250 | 500 | 250 |
| 4.00 | 100 | 100 | 100 | 100 | 100 | 100 | 100 | 250 | 100 |
| 4.20 | 100 | 100 | – | 100 | 100 | 100 | 100 | 100 | 100 |
| 4.40 | 100 | 100 | – | 100 | 100 | 100 | 100 | 100 | 100 |
| 4.60 | – | 100 | – | 100 | 100 | 100 | 100 | 100 | 100 |
| 4.80 | – | – | – | 100 | 100 | 100 | 100 | 100 | 100 |
| 5.00 | – | – | – | 100 | 100 | 100 | 100 | 100 | 100 |
| 5.20 | – | – | – | – | 100 | 100 | – | 100 | 100 |
| 5.40 | – | – | – | – | 100 | 100 | – | 100 | 100 |
| 5.60 | – | – | – | – | – | 100 | – | 100 | 100 |

To obtain smooth and well-converged PMFs, careful choice of umbrella window force constants is required. Table S1 shows the force constants used to obtain the PMFs (Figure 1) after sometimes as much refinement as 3 rounds, for the systems of POPC 100, DOPE:DOPS:DOPC 60:20:20 (DDD), and DOPE:DOPS:DOPC:Chol 40:20:20:20 (DDDC).

#### Which Lipids Peptides Prefer

For each umbrella window, the radial distribution function of lipids around the center of mass (COM) of the peptide in the $x$-$y$ membrane plane was calculated. See Figure S1 and Figure S2 for the membrane compositions without and with cholesterol, respectively.

On the $x$ and $y$ axes are the membrane-peptide COM-COM $z$-distance of separate umbrella window simulations and peptide-lipid distances in the $x$-$y$ membrane plane, respectively. The $z$ axis shows the value of the radial distribution function. If the RDF is below the value 1, representing the ideal lipid mixing in infinite peptide-lipid $x$-$y$ plane distances, the value is colored blue. On the other hand, values above 1, representing excessive lipid occurrence, are colored red.

#### Force Field Comparison

Due to the overestimation of interactions involving charged species by traditional, non-polarisable force fields, most of the simulations performed here used the ProsECCo force field.[S1]

For comparison, figure S3 shows the calculated free energy profiles for $R_4$, $K_9$, and $R_9$ binding to the DDDC membrane calculated from simulations using ProsECCo, as well as the $R_9$ free energy profile obtained using the CHARMM force field.

The free energy minimum of this interaction is much stronger than in the case of ProsECCo, $-78.67 \pm 7.61$ kJ/mol. Similarly to ProsECCo results, the CHARMM $R_9$ PMF minimum is located at the 2.35 nm peptide-membrane center of mass $z$-distance.

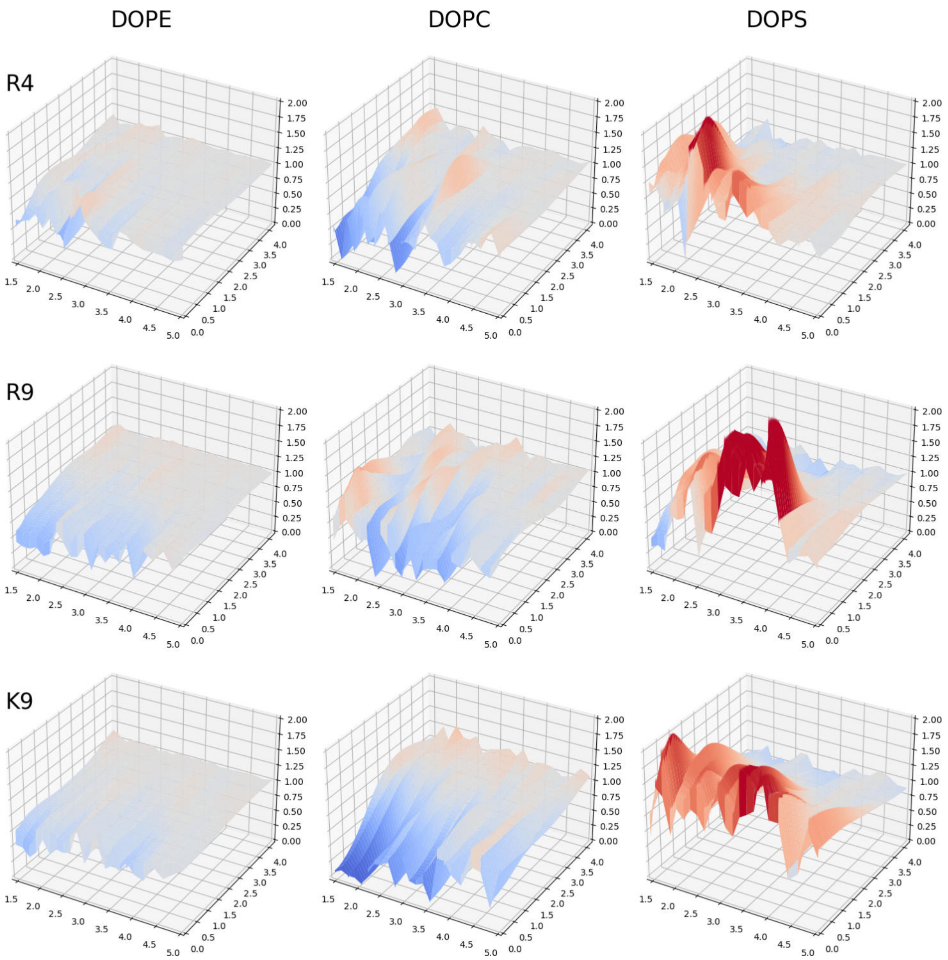


Figure S1: **Lipid density maps for DDD system** shows a planar radial distribution function of lipids around the peptide in the membrane $x$-$y$ plane for selected umbrella windows. In the $x$ axis, each umbrella window is represented by a peptide-membrane center of mass $z$-distance $d$ (in other terms, how high is the peptide above the membrane). The $y$ axis represents peptide-lipid distance in a plane and, finally, the $z$ axis is the value of the radial distribution function in the lipid membrane plane. The blue color reflects RDF values less than 1, indicating a lipid deficiency in that region, whereas the red color points to the excessive presence of the particular lipid in said region, both compared to the average value at infinite distance.

First row depicts simulations containing tetraarginine, namely DOPE, DOPC and DOPS lipids around $R_4$ on left, middle and right, respectively. The second and third row similarly shows peptide-lipid RDFs for nonaarginine and nonalysine.

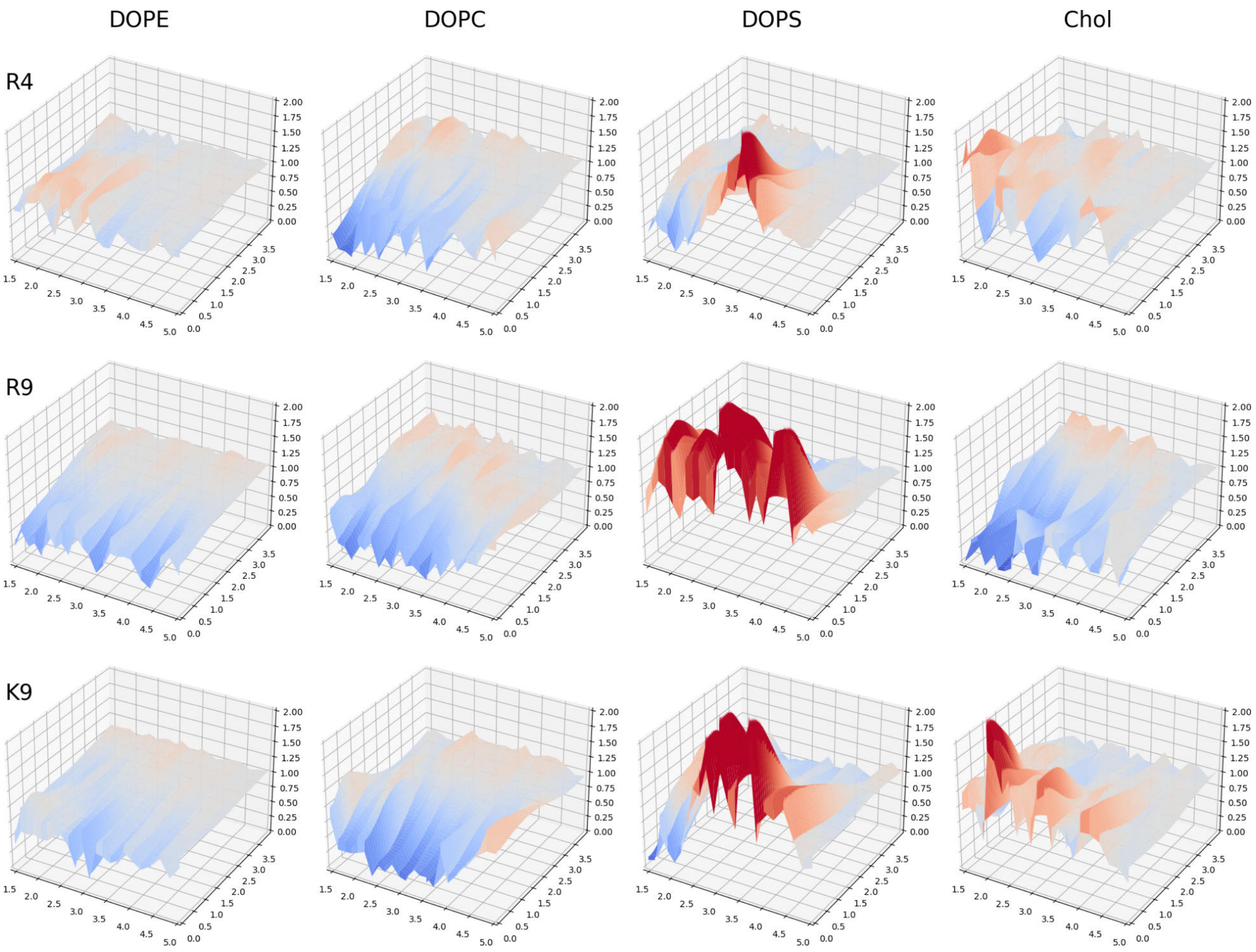


Figure S2: **Lipid density maps for DDDC system** shows a planar radial distribution function of lipids around the peptide in the membrane $x$-$y$ plane for selected umbrella windows.

Similarly as in Figure S1, $x$ axis represents the umbrella window distance $d$, meaning how high is the peptide COM in $z$ direction above membrane COM. Axis $y$ is the peptide-lipid distance in an $x$-$y$ membrane plane and $z$ is the RDF value, coloured blue and red when lower and higher than 1, respectively.

Top row corresponds to simulations containing $R_4$, middle and bottom raw $R_9$ and $K_9$, respectively. The first column shows peptide-DOPE RDFs, the second column depicts RDF between peptide and DOPC lipids, peptide-DOPS RDFs are in the third column, and cholesterol radial distribution functions around peptide are in the fourth column.

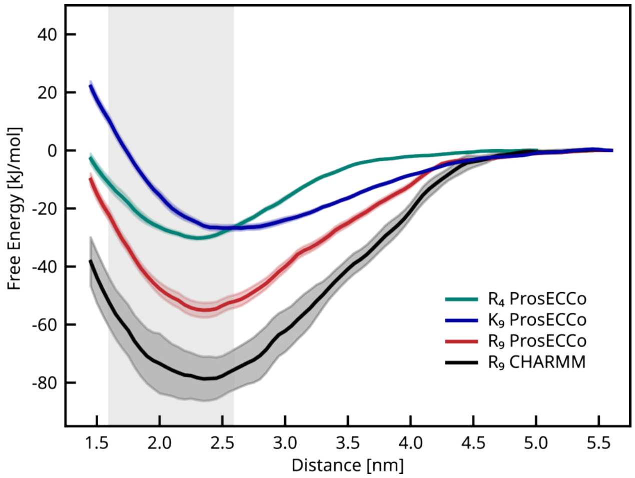


Figure S3: **Free energy profiles of peptide interactions with a DDDC membrane.** Grey area indicate lipid headgroup region.

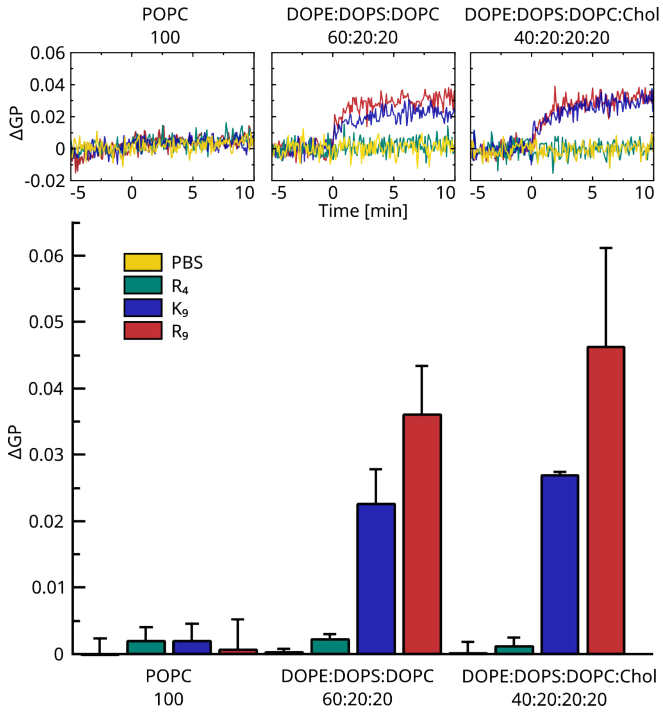


Figure S4: **Peptide effect on various lipid membranes** depicted as changes of a general polarisation factor (ΔGP) of laurdan spectra after adding $R_4$ (green), $K_9$ (blue) or $R_9$ (red) peptides or no peptide (PBS +/+ buffer control; yellow) to the LUV solution at time 0 min **in the presence of $Ca^{2+}$ and $Mg^{2+}$ ions**. Top: Examples of ΔGP time evolution for three different LUV membrane compositions: POPC (left), DDD (middle), and DDDC (right). Bottom: The average GP increase for three independent measurements after peptide addition.

### Peptide Effects on Membranes Quantified by Laurdan GP Changes

#### Calcium-Containing LUV Suspensions

To investigate a possible effect of $Ca^{2+}$ and $Mg^{2+}$ ions on peptide-membrane interactions measured as a change of membrane-embedded fluorescent dye Laurdan's GP, the main article LUV experiments were replicated using calcium- and magnesium-containing buffer PBS +/+. These experiments were done in triplicate and the average GP changes with errors are in Figure S4, resulting in similar trends as observed in systems lacking $Ca^{2+}$ and $Mg^{2+}$ ions.

#### OVCAR-3 Extracellular Vesicles

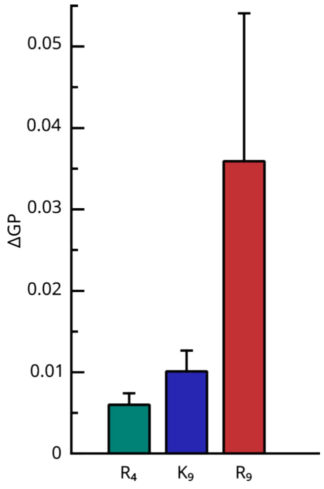


Figure S5: **Peptide effect on OVCAR-3 extracellular vesicle membranes** depicted as changes of a general polarisation factor (ΔGP) of laurdan spectra after adding $R_4$ (green), $K_9$ (blue) or $R_9$ (red) peptides to the EV solution. The average GP increase for three independent measurements after peptide addition.

Embedding laurdan into the membranes of extracellular vesicles extracted from the ovarian cancer cell line OVCAR-3 enables us to measure the shift in laurdan's spectra, quantified by ΔGP. Similarly to the case of lipid-only LUVs, the biggest effect on EV membranes has nonaarginine; see Figure S5. The addition of nonalysine results in a very low GP change, only slightly higher than that of tetraarginine. All experiments were performed in triplicate.

### I. Monte Carlo Simulations

Uses `organl` in a slight modification of[S2] The computational and analysis methodology also follows the same framework as described in our previous work[S2] where a detailed description of the non-interacting benchmark used in this study is provided.

The interactions between the surface elements are modeled using a Lennard-Jones 12–6 potential of the form:

$$dU = 4\epsilon \left[ \left( \frac{\sigma}{r_{12}} \right)^{12} - \left( \frac{\sigma}{r_{12}} \right)^{6} \right] dA_1 \, dA_2, \tag{1}$$

where $r_{12}$ is the distance between differential surface elements $dA_1$ and $dA_2$. The potential exhibits a minimum at $r_{\text{min}} = 2^{1/6}\sigma$ nm, with a corresponding energy value of $U = -\epsilon\,\text{kT/nm}^2$. In this study, we fix $r_{\text{min}} = 5$ nm. Interactions are evaluated only when both participating surface elements contain $R_9$ and the angle between their respective surface normals exceeds 30°. Surface integrals are computed using a 7-point symmetric Gauss quadrature rule defined over triangular elements.

##### Simulation Details

Simulations were conducted for over 185000 Monte Carlo steps across a range of $\epsilon$ values. It was observed that for $\epsilon > 0.012$ kT/nm$^2$, corresponding to a potential minimum exceeding 2.4 kJ/mol, the invaginated morphology deviated significantly from the reference shape obtained without interactions. These deviations manifested as pronounced elongation or pancaking of the invagination.

**Energy profiles** are computed by averaging over every 500 datapoint spread across the entire simulation runthe final 500 outputs, corresponding to the last 185000 Monte Carlo steps. The error bars reported represent the 95% Confidence Interval of standard error on the mean corrected for temporal correlations inherent in Monte Carlo sampling using the statistical inefficiency estimate.[S3]

**Bin Averaging** for Fig. S6 and S7: To preserve spatial correlations for curvature analysis, we applied a bin-averaging method to the snapshots. Local properties (e.g., lipid density) were sorted into bins based on a second variable (e.g., mean curvature) and averaged within each bin across all snapshots. This approach maintains the relationship between geometric features and local composition without losing detail through spatial averaging.

#### Results

A comparison of Figures S6 and S7 indicates that, in the presence of interactions, the majority of $R_9$ predominantly localize within the neck (region B) and the invagination (region C). Notably, no discernible changes in lipid patterning relative to the $R_9$ distribution are observed under either condition.

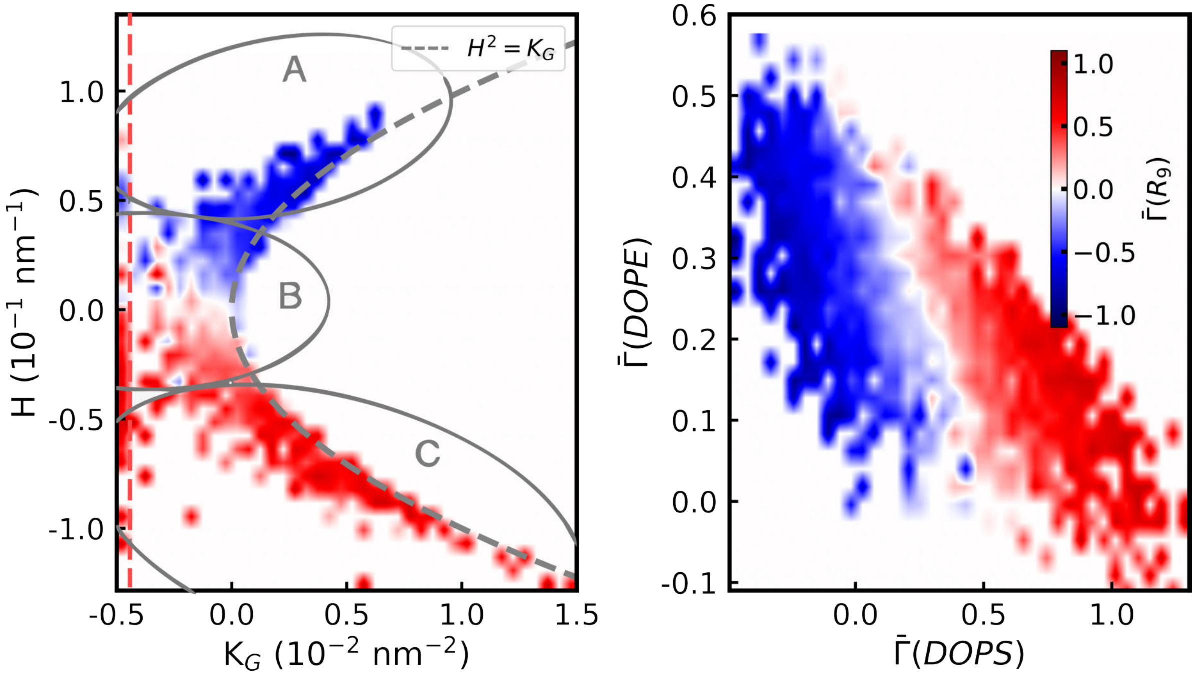


Figure S6: **Sorting in the Absence of Interaction Potential.** (Left) Bin-averaged heatmap of excess $R_9$ coverage on a stomatocyte at a reduced volume $\nu_0 = 0.90$. Regions A, B, and C correspond to the exterior, neck, and invagination, respectively. The dashed line indicates excluded regions, where outliers were removed to enhance visual clarity. (Right) Heatmap of excess $R_9$ coverage as a function of DOPE and DOPS composition. These data also constitute a reproduction of what was reported in,[S2] Fig. 7

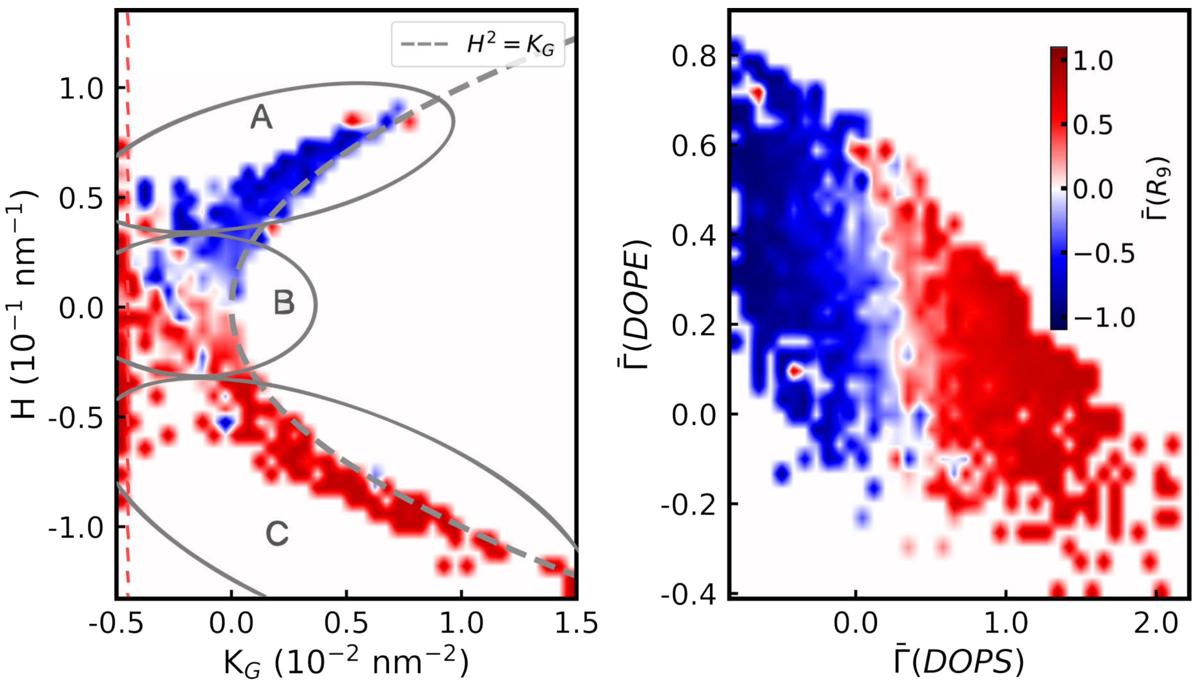


Figure S7: **Sorting in Presence of Interaction Potential.** (Left) Bin-averaged heatmap of excess $R_9$ coverage on a stomatocyte at a reduced volume $\nu_0 = 0.90$. Regions A, B, and C correspond to the exterior, neck, and invagination, respectively. The dashed line indicates excluded regions, where outliers were removed to enhance visual clarity. (Right) Heatmap of excess $R_9$ coverage as a function of DOPE and DOPS composition.

### II. Continuum Model for Vesicle Inbudding Phase Diagram

We briefly describe the continuum model used in this study, which extends a previously developed model[S4] by incorporating multicomponent membranes, a peptide field on the outer leaflet, and neck adhesion mediated by the $R_9$ peptide. Building on the Helfrich–Hamm–Kozlov framework[S5–S7] and its extensions by Ryham et al.,[S8–S10] the model describes bilayer vesicles with cylindrical symmetry.

Each bilayer leaflet ($j = o, i$ for outer and inner) is represented by a lipid vector field $\boldsymbol{D}_j = l_j \boldsymbol{n}_j$, where $l_j$ is the relaxed lipid length and $\boldsymbol{n}_j$ the unit director. This field is defined on the leaflet's neutral surface ($S_j$) and is oriented toward the midplane. The elastic contribution to the free energy of each leaflet includes contributions from bending, tilt, and area compressibility:

$$F_j^{\text{el}} = \frac{1}{2}\int_{S_j} \left[K_C\left((2H+J_s)^2 - J_s^2\right) + \kappa_t \boldsymbol{t}^2 + K_a\alpha^2\right] \mathrm{d}S_j \tag{2}$$

The first term represents the bending energy, with the bending modulus $K_C$ determining the energy penalty for splay deformations, where $2H = \nabla \cdot \boldsymbol{n}$, and the natural curvature is given by the spontaneous curvature $J_s$ (positive for a spherical micelle). The second term describes the tilt energy governed by the tilt modulus $\kappa_t$ and the tilt deformation $\boldsymbol{t} = \boldsymbol{n}/(\boldsymbol{N}\cdot\boldsymbol{n}) - \boldsymbol{N}$, where $\boldsymbol{N}$ is the surface normal. The final term defines the area-compressibility energy with modulus $K_a$ for deviations in the local area-per-lipid (APL), $a$, from its relaxed value $a_0$, described by $\alpha = \frac{a-a_0}{\sqrt{a_0 a}}$. Under the assumption of local lipid incompressibility ($al = a_0 l_0$), this becomes $\alpha = \frac{l_0 - l}{\sqrt{l_0 l}}$.

Volume constraints enforce lipid incompressibility and maintain a constant entrapped water volume, consistent with vesicle formation physics and experimental observations.[S11–S13]

For systems with $1 < m$ lipids, an additional $m$-vector field $\boldsymbol{\phi} = (\phi_0, \ldots, \phi_{m-1})$ is defined on the leaflets' surface; this field accounts for the molar ratio of each lipid. The constrained nature of the molar ratios, i.e. $\phi_i \in (0,1)$ and they must sum to 1, can complicate numerical minimization methods. To address this, an unconstrained $(m-1)$-vector field $\boldsymbol{\Phi}_+$ is introduced. The actual molar composition $\boldsymbol{\phi}$ is obtained recursively as:

$$\begin{aligned}
\phi_1 &= 0.5 \cdot (\tanh(\Phi_{+,1}) + 1) \\
\phi_2 &= (1-\phi_1) \cdot 0.5 \cdot (\tanh(\Phi_{+,2}) + 1) \\
\phi_3 &= (1-\phi_1)(1-\phi_2) \cdot 0.5 \cdot (\tanh(\Phi_{+,3}) + 1) \\
&\vdots \\
\phi_{m-1} &= \left[\prod_{i=1}^{m-2}(1-\phi_i)\right] \cdot 0.5 \cdot (\tanh(\Phi_{+,m-1}) + 1)
\end{aligned}$$

with $\phi_0$ deduced by $1 - \sum_{i=1}^{m-1}\phi_i$. This transformation ensures that all components lie within

(0, 1) and sum up to 1, enabling stable numerical optimization.

Lipid mixing is accounted for by the additional energy density contribution:

$$f^{\text{mix}}(\phi) = \left( \sum_{i=0}^{m-1} \phi_i \log\left(\frac{\phi_i}{\phi_{0,i}}\right) + K \sum_{i=1}^{m-1} |\nabla \phi_i|^2 \right) \frac{k_{\text{B}}T}{a} \tag{3}$$

The first term corresponds to the ideal mixing entropy of lipids over the surface, where $\phi_{0,i}$ denotes the average molar fraction of the $i$-th lipid component. In this study, we used a ternary lipid composition with mean molar fractions $\phi_{\text{DOPE}} = 0.6$, $\phi_{\text{DOPS}} = 0.2$, and $\phi_{\text{DOPC}} = 0.2$. The second term enforces the smoothness of $\phi$ across the surface. The parameter $K$ is set to a small value of $0.2\,\text{nm}^2$, ensuring minimal impact on the results. This choice corresponds to a line tension of approximately $0.25\,k_{\text{B}}T\,\text{nm}^{-1}$ for a linear gradient of $\phi$ from 0 to 1 over 0.8 nm, roughly the lateral length scale of a lipid, consistent with line tensions observed in liquid-liquid phase coexistence.[S14] This selection guarantees that lipid domains—regions with significantly high or low lipid concentration—are at least nanometric in size, exceeding both the model resolution and typical lipid lateral dimensions. When there are no domains, $|\nabla \phi_i|^2 \ll 0.01\,\text{nm}^{-2}$, leading to negligible energy contributions and minimal composition variance. Choosing significantly smaller values of $K$ risks discontinuities in $\phi$ due to the finite model resolution, whereas larger values may inhibit domain formation by penalizing interfaces, thus affecting the results. Similarly to the volume constraint, compositional constraints are imposed to maintain a fixed total lipid concentration. Furthermore, the local spontaneous curvature is expressed as a linear combination of the components:

$$J_s = \sum_{i=0}^{m-1} \phi_i \, J_{s,i}$$

where $J_{s,i}$ is the spontaneous curvature of the $i$-th component.

For systems containing $R_9$ on the outer leaflet, a scalar field $\phi_p$ is introduced to represent the local occupancy of $R_9$. It is defined and implemented analogously to the $\boldsymbol{\phi}$ field, but with $m = 1$. The corresponding mixing energy density is given by:

$$f^{\text{mix}}(\phi_p) = \left(\phi_p \log\left(\frac{\phi_p}{\phi_{p,0}}\right) + (1-\phi_p)\log\left(\frac{1-\phi_p}{1-\phi_{p,0}}\right)\right)\frac{k_{\text{B}}T}{A_{\text{R}_9}} + K\left|\nabla\phi_p\right|^2\frac{k_{\text{B}}T}{a} \tag{4}$$

where $A_{\text{R}_9} = 12.31$ nm is the area per $\text{R}_9$ site on the membrane,[S2] and $\phi_{p,0}$ is the enforced mean occupancy of $\text{R}_9$. The local spontaneous curvature is then defined as a weighted linear combination of the spontaneous curvatures for membranes with and without $\text{R}_9$:

$$J_s = (1-\phi_p)\, J_s^{\text{no R}_9} + \phi_p\, J_s^{\text{with R}_9}.$$

Notably, both $J_s^{\text{with R}_9}$ and $J_s^{\text{no R}_9}$ are themselves linearly interpolated across lipid components in systems containing multiple lipids.

When incorporating neck adhesion mediated by $\text{R}_9$ binding, we add an adhesion free energy density term:

$$f_{\text{ad}}(r,\theta) = -\phi_p \cdot \epsilon \cdot g\big(h(r)\big) \cdot g\big(k(\theta)\big) \tag{5}$$

where $\epsilon$ denotes the binding energy density at the neck surface for radius $R$. Adhesion is activated smoothly according to both radial and angular criteria: it occurs when the angle $\theta$ between the outer leaflet normal and the $-\hat{\mathbf{r}}$ direction lies between 30° and 150°, and when the local radius $r$ is near $R + l$ (set to 2 nm), reaching its maximum at $r = R$ (set to 1 nm). These values are consistent with the characteristic size of $\text{R}_9$ peptides. The activation function $g(x)$ is defined piecewise as

$$g(x) = \begin{cases} 1 & x \le -0.5 \\ 2x^3 - 1.5x + 0.5 & -0.5 < x < 0.5 \\ 0 & x \ge 0.5 \end{cases}$$

ensuring a smooth transition in activation. The angular dependence is captured by

$$k(\theta) = \begin{cases} -2\sin(\theta) + 1.5 & 30^\circ < \theta < 150^\circ \\ 0 & \text{otherwise} \end{cases}$$

and the radial dependence by

$$h(r) = \frac{r - (R - l)}{R} - 0.5$$

defined over the activation region. The binding energy density, $\epsilon$, was varied across different systems. In most cases, it was set to 0. In the "Sticky Neck" simulations, it was set to either $6\,\mathrm{pN} \cdot \mathrm{nm}^{-1}$ or $12\,\mathrm{pN} \cdot \mathrm{nm}^{-1}$. The value of $6\,\mathrm{pN} \cdot \mathrm{nm}^{-1}$ corresponds roughly to half the binding energy of a single $R_9$ peptide to a DOPS membrane,[S2] divided by the area per $R_9$, $A_{R_9}$. This estimate is reasonable under the assumption that $R_9$-membrane interactions are additive in the cylindrical neck region.

For a vesicle with midplane radius $R_m$, the equilibrium phase is determined by the leaflet area asymmetry ($\Delta A$) and the relative water content ($\nu$). The asymmetry is given by $\Delta A = 4l_0 R_m m_0$, where $m_0$ quantifies the intrinsic area mismatch between the leaflets. The entrapped water volume is expressed as $V_w = \nu V_w^{\mathrm{id}}$, with the ideal volume defined by $V_w^{\mathrm{id}} = \frac{4\pi}{3}(R_m - l_0)^3$. To determine which morphology is energetically favorable under given conditions, we calculate the total energy of the vesicle in both the inbudded and unbudded states. This comparison allows us to identify the phase boundary where the two configurations coexist, i.e., where their energies are equal. Mapping this boundary across different values of $m_0$ and $\nu$ produces the phase diagram. For further details on the methodology used to compute phase diagram coexistence lines or structural optimization, please refer to the Supporting Information of Ref. S4.

We ran a hierarchy of models with increasing levels of biophysical detail to systematically assess the impact of each factor on the vesicle inbudding tendency. In all simulations, we fixed the midplane vesicle radius to $R_m = 50\,\mathrm{nm}$, lipid length to $l_0 = 1.55\,\mathrm{nm}$, and relaxed area-per-lipid to $a_0 = 0.625\,\mathrm{nm}^2$, with the system held at temperature $T = 298.15\,\mathrm{K}$. Elastic parameters were set to $K_C = 15\,k_\mathrm{B}T$, $\kappa_t = 10\,k_\mathrm{B}T \cdot \mathrm{nm}^{-1}$, and $K_a = 55\,k_\mathrm{B}T \cdot \mathrm{nm}^{-1}$. These parameters closely match those observed in simulations (see Ref. S2 and the Complementary Simulations subsection), while ensuring numerical stability and consistency with continuum-level behavior. The spontaneous curvature values were directly taken from molecular dynamics simulations in Ref. S2. In single-component systems, effective spontaneous curvature was applied directly, with $J_s = -0.156\,\mathrm{nm}^{-1}$ for membranes without $\mathrm{R}_9$ ($\phi_p = 0$) and $J_s = -0.214\,\mathrm{nm}^{-1}$ for full $\mathrm{R}_9$ occupancy ($\phi_p = 1$). For multicomponent systems, the component-specific values were as follows: DOPE, $J_s = -0.240\,\mathrm{nm}^{-1}$ without $\mathrm{R}_9$ and $J_s = -0.307\,\mathrm{nm}^{-1}$ with $\mathrm{R}_9$; DOPC, $J_s = 0$ without $\mathrm{R}_9$ and $J_s = -0.043\,\mathrm{nm}^{-1}$ with $\mathrm{R}_9$; DOPS, $J_s = -0.060\,\mathrm{nm}^{-1}$ without $\mathrm{R}_9$ and $J_s = -0.104\,\mathrm{nm}^{-1}$ with $\mathrm{R}_9$. In simulations that included the peptide, we fixed the mean occupancy to $\phi_p = 0.5$.

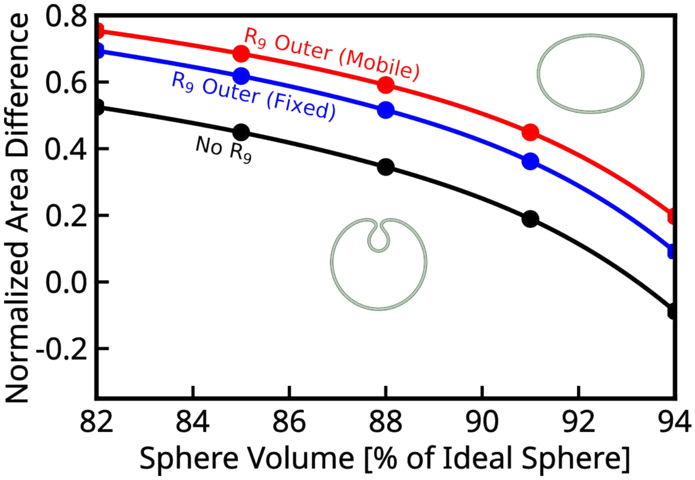


Figure S8: **Phase diagram of the continuum model** for multi-component systems in the $(\nu, m_0)$ parameter space, following the inbudding transition. The coexistence curve (lines), interpolated with a cubic spline, separates the oblate and inbudded states from above and below, respectively, and is derived from sampled data points (circles). The diagram reflects variations in conditions such as the presence of $\mathrm{R}_9$ on the outer leaflet ($\mathrm{R}_9$ Outer) and its redistribution across the membrane ($\mathrm{R}_9$ Outer, Mobile).

#### Complementary

To determine appropriate values for the area per lipid ($a_0$), leaflet thickness ($l_0$), and area compressibility modulus ($K_a$) in the model, we performed complementary all-atom MD simulations. The systems consisted of DOPE:DOPS:DOPC (2:1:1) membranes with 400 lipids and varying amounts of $R_9$, constructed using CHARMM-GUI[S15] and the ProsECCo force field.[S1] The thickness of the water layer was set to 15 nm, with NaCl at an ionic concentration of 150 mM. The $R_9$ peptides were initially placed symmetrically near both leaflets (i.e., half near the upper leaflet and half near the lower leaflet). The temperature was maintained at 298 K. Equilibration and production followed the standard CHARMM-GUI protocol. Each simulation was run for 300 ns, with the first 100 ns treated as equilibration and excluded from the analysis. The results of the analysis are given in Table S2.

For each system, the relevant properties were calculated as follows. The area per lipid was obtained by dividing the average lateral ($xy$) simulation box area by the number of lipids per leaflet. The leaflet thickness was estimated by calculating the average $z$-position of the $C_1$ atoms in each leaflet, which approximates the neutral plane height.[S16] The average distance between these planes was computed and halved to obtain the average leaflet thickness. The area compressibility modulus, $K_a$, for planar bilayers was estimated from area fluctuations in periodic bilayer simulations[S17] using

$$K_a = \frac{A_0}{2\,\mathrm{Var}(A)}\, k_{\mathrm{b}}T\,\mathrm{nm}^{-1},$$

where $A$ is the instantaneous monolayer lateral area. Standard errors were estimated by 10-bins block averaging.

Table S2: **Complementary MD simulations results** - DOPE:DOPS:DOPC (2:1:1) membranes with varying $R_9$ content. Values are averages over the last 200 ns of production simulations.

| $R_9$ content | $a_0$ [nm$^2$] | $l_0$ [nm] | $K_a$ [k$_B$T] |
|---|---|---|---|
| 0 $R_9$ | 0.6262±0.0006 | 1.564±0.001 | 52±8 |
| 8 $R_9$ | 0.6249±0.0007 | 1.565±0.002 | 65±10 |
| 16 $R_9$ | 0.626±0.001 | 1.563±0.003 | 59±14 |
| 32 $R_9$ | 0.625±0.001 | 1.563±0.003 | 55±10 |

### Cryo-TEM and Cryo-ET

#### Cryo-TEM

Cryo-TEM visualization of LUVs of differing lipid composition (POPC S9 A, DDD S9 B, and DDDC S9 C), both with all the tested peptides and without peptide. We clearly see that POPC-only liposomes show no membrane remodelling, even $R_9$ induces only vesicle-vesicle aggregation and adhesion, without forming multilamellar structures. Shifting the membrane composition to DOPE- and DOPS-rich, we see a sharp difference. For DDD LUVs, longer peptides induce significant remodelling, with $K_9$ resulting in occasional bifurcation and $R_9$ displaying very strong multilamellarity due to high vesicle-vesicle interactions. Conversely, for DDDC LUVs, the effect of $K_9$ is much more limited and $R_9$ results in clear multilamellarity instead. In both PE-containing compositions, incubation with $R_4$ does not result in membrane remodelling.

#### Extracellular Vesicles

To further understand how membrane composition may affect the $R_9$ interaction with membranes, we performed cryo-ET visualization of extracellular vesicles (EVs) - which fully retain the cell plasma membrane compositional complexity, including protein and glycans - incubated for 10 min with 150 $\mu$M $R_9$ (Figure S9). Similarly to $R_9$ interaction with lipid-only vesicles, we observe membrane remodelling in the form of bilamellar morphologies (Fig. S10, A), alongside unaltered EVs (Figure S10, B) or cases in which vesicles appear burst (Fig-

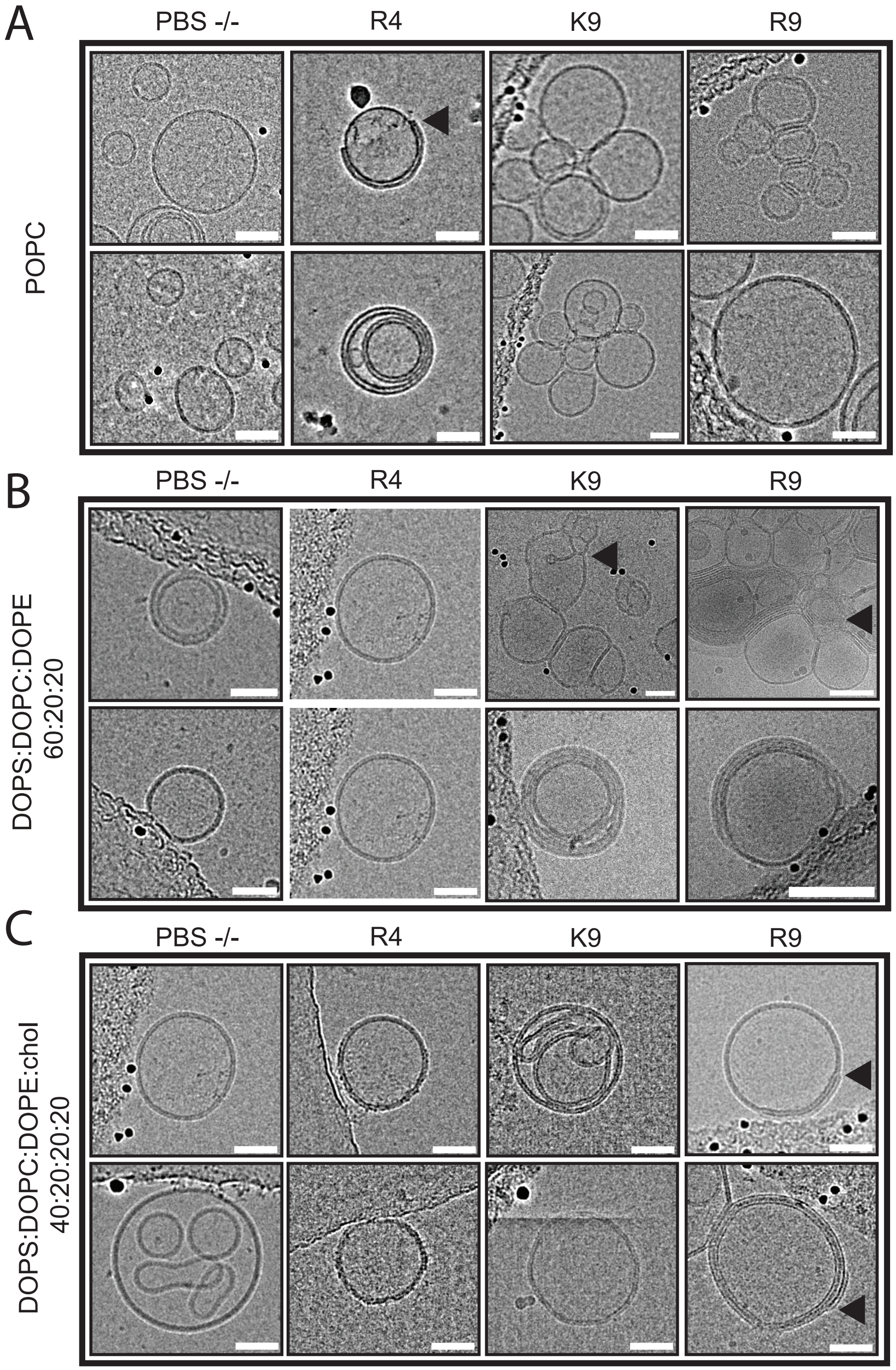


Figure S9: **Examples of cryoTEM images of LUVs** composed of either POPC (A), DDD (B), and DDDC (C) interacting with either no peptides, $R_9$, $K_9$ or $R_9$. Scale bar is 50 nm.

ure S10, C).

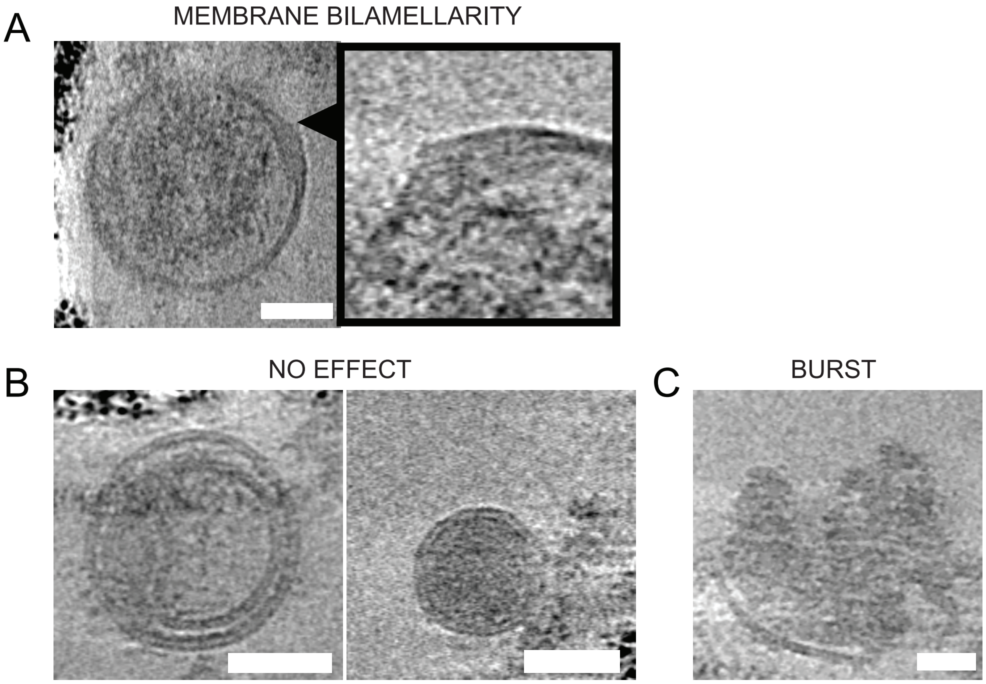


Figure S10: **Representative cryo-ET images of EVs.** (A) EVs incubated with $R_9$ for 10 min resulting in clear membrane bilamellarity (blue triangles) with zoomed-in detail on the remodelling spot (blue squares, scale bar: 20 nm). (B) EVs incubated with $R_9$ showing no clear morphological change or remodelling. (C) EV broken upon incubation with $R_9$. In all images except details, the scale bar is 50 nm.

#### Selected Examples

Figure S11, A, shows vesicles containing inner membrane structures (left, without $R_9$) and examples of structures observed with various $R_9$ incubation times. We hypothesise that, with time, nonaarginine gradually depletes the vesicle inner membrane structure, resulting in substantial multilamellarity, as visible in the 10 min examples. Also note that at 10 min, dark spots indicating $R_9$ are present inside the vesicles.

Inward budding structures captured by cryo-ET are shown in Figure S11, B. This $R_9$-induced membrane remodelling is limited, possibly due to the lack of an available membrane reservoir, compared to the extensive membrane remodelling of PS-rich vesicles rich in inner

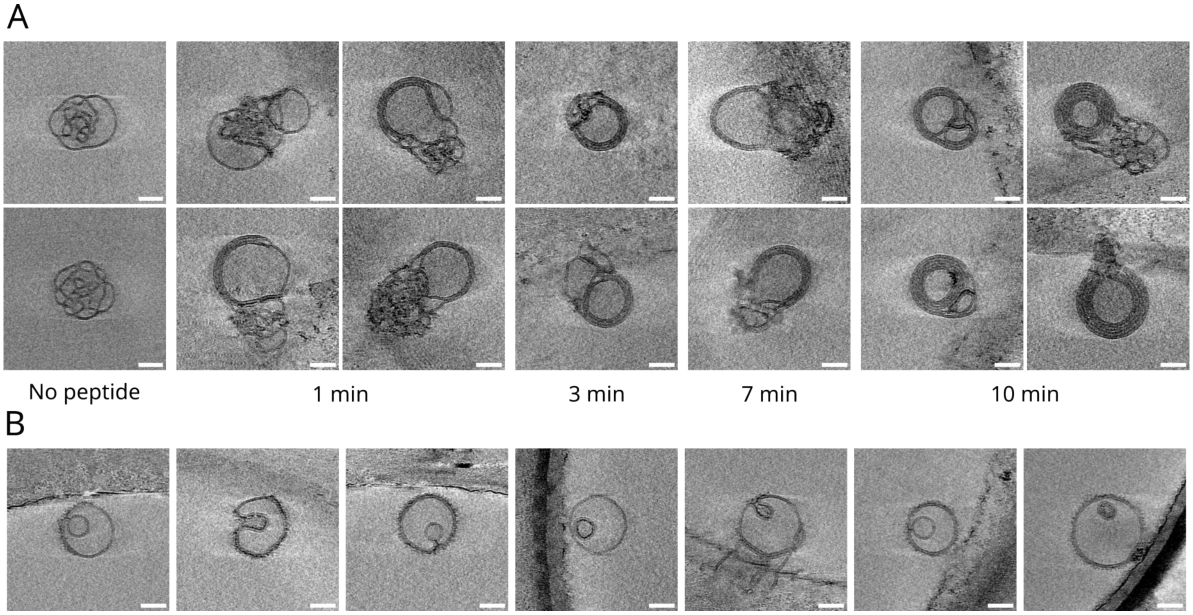


Figure S11: **Structures observed by cryo-electon tomography.** (A) Proposed time evolution of a PS-rich LUV subpopulation ($\sim$ 15%). On the left, two PS-rich vesicles without nonaarginine presence are shown. PS-rich vesicles incubated with $R_9$ for 1–10 min are gradually disentangled, their inner membrane sponge-like mesh transforming into the final multilamellar structures. Note that darker regions between membranes as well as on the inner vesicle side at 10 min timepoint is nonaarginine. (B) Examples of observed membrane budding structures. $R_9$ is visible on the vesicle membrane or in the invaginations structure as a fluffy dark membrane cover. If only a circle is visible inside the LUV vesicle, the invagination is occuring in the $z$-plane (either from top or the bottom of the vesicle, not sideways). Scale bar is 50 nm.

membrane structure (shown in Figure S11, A).

#### Neural Network Training

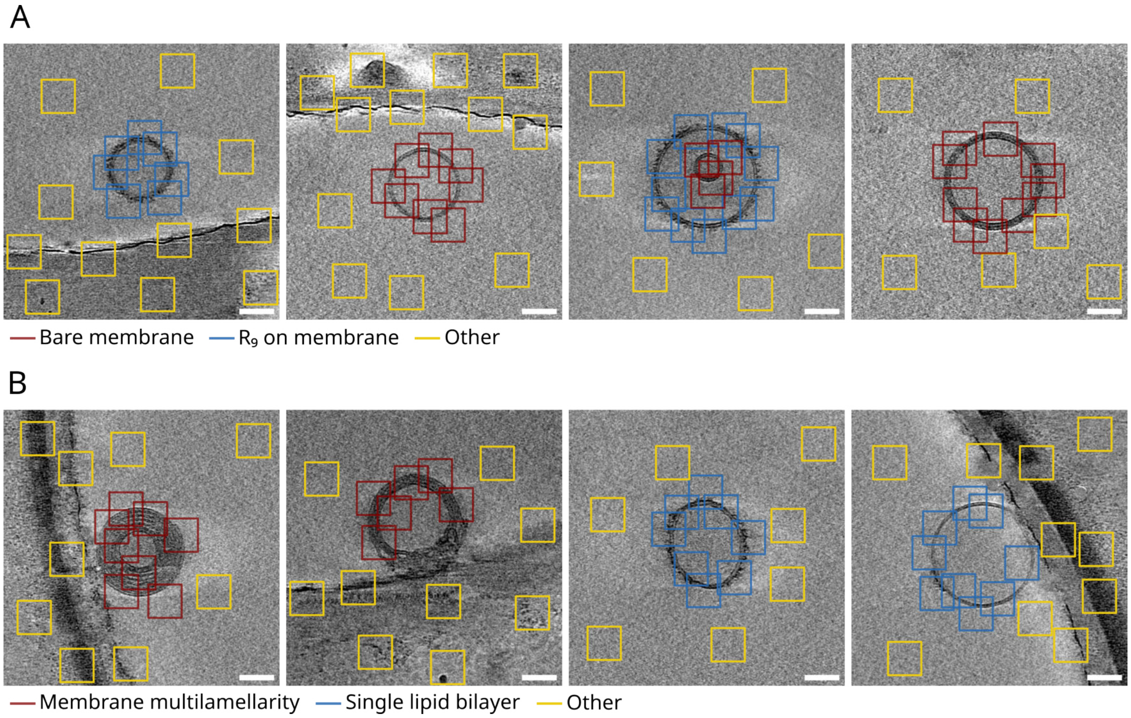


Figure S12: **Neural network training examples.** (A) Training to distinguish $R_9$ peptide on the the outer vesicle membrane (blue) from lipid membrane without nonaarginine (red) and the outer space, including the EM grid edge (yellow). (B) Unilamellar (blue) versus multilamellar (red) membrane training examples. $R_9$ on a single membrane is considered unilamellar vesicle part. Again, non-vesicle training parts are shown yellow. Scale bar is 50 nm.

Figure S12 shows example snapshots from dataset training for both "$R_9$ on the outer membrane" and "Multilamellarity" feature training - manually selecting features of interest in 64x64 pixel squares.

#### Cell Translocation Depended on Peptide Concentration

To test the dependence of cell translocation on the $R_9$ concentration, U-2 OS cells were incubated and imaged with 1–15 $\mu$M nonaarginine (see Figure S13) for 20 min.

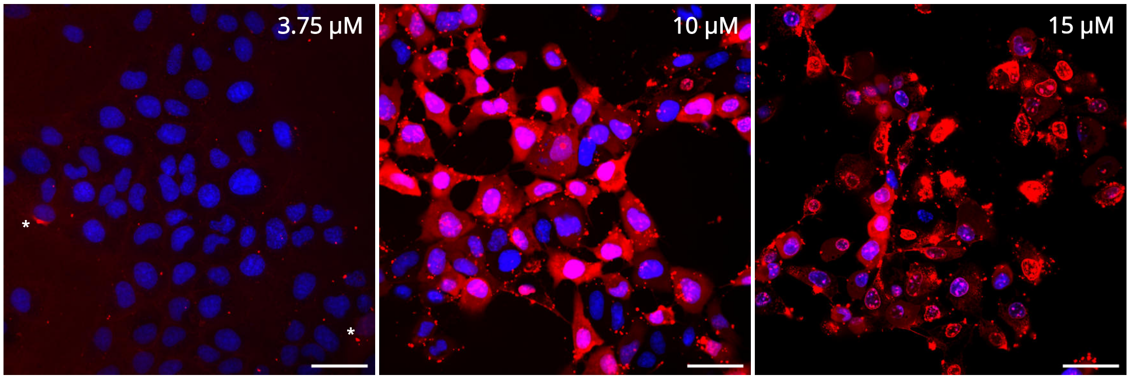


Figure S13: **Cell penetration depending on $R_9$ concentration.** Cells were incubated with Tamra-labelled $R_9$ (red) for 20 min. Cell nuclei is stained by Hoechst dye (blue). Left: Cells incubated with 3.75 $\mu$M nonaarginine. White asterisks show two passively penetrated cells. Middle: Cells incubated with 10 $\mu$M $R_9$. Right: Rapid cell translocation and death is observed with 15 $\mu$M $R_9$ incubation. Scalebar: 50 $\mu$m.

We observed that below approximately 3.75 $\mu$M of $R_9$ concentration, cells rarely get penetrated passively. Rather, small, endocytic-like dots are observed. Figure S13, left, shows a threshold situation, with two cells passively penetrated by $R_9$ (emphasized by white asterisks) and a few $R_9$-membrane puncta that we connect to the passive cell penetration. Much smaller, endocytic-like dots, are abundant.

A passive cell penetration is shown in Figure S13, middle and right. Cell translocation is so rapid at 15 $\mu$M $R_9$ concentration, that most cells burst and die (Figure S13, right).

### Flow Cytometry

#### Concentration Study

As a data processing example, 15 $\mu$M peptide concentration measurements are shown in Figure S14. In the first gating step (first row of Figure S14, A), only viable cells-events were selected. Among these, the second gating selects singlet cells-events from the previously selected (second row). Lastly, a fluorescence intensity histogram was calculated for this selection (third row). The columns represent, from left to right, cells control without peptide presence, $R_4$, $K_9$, and $R_9$.

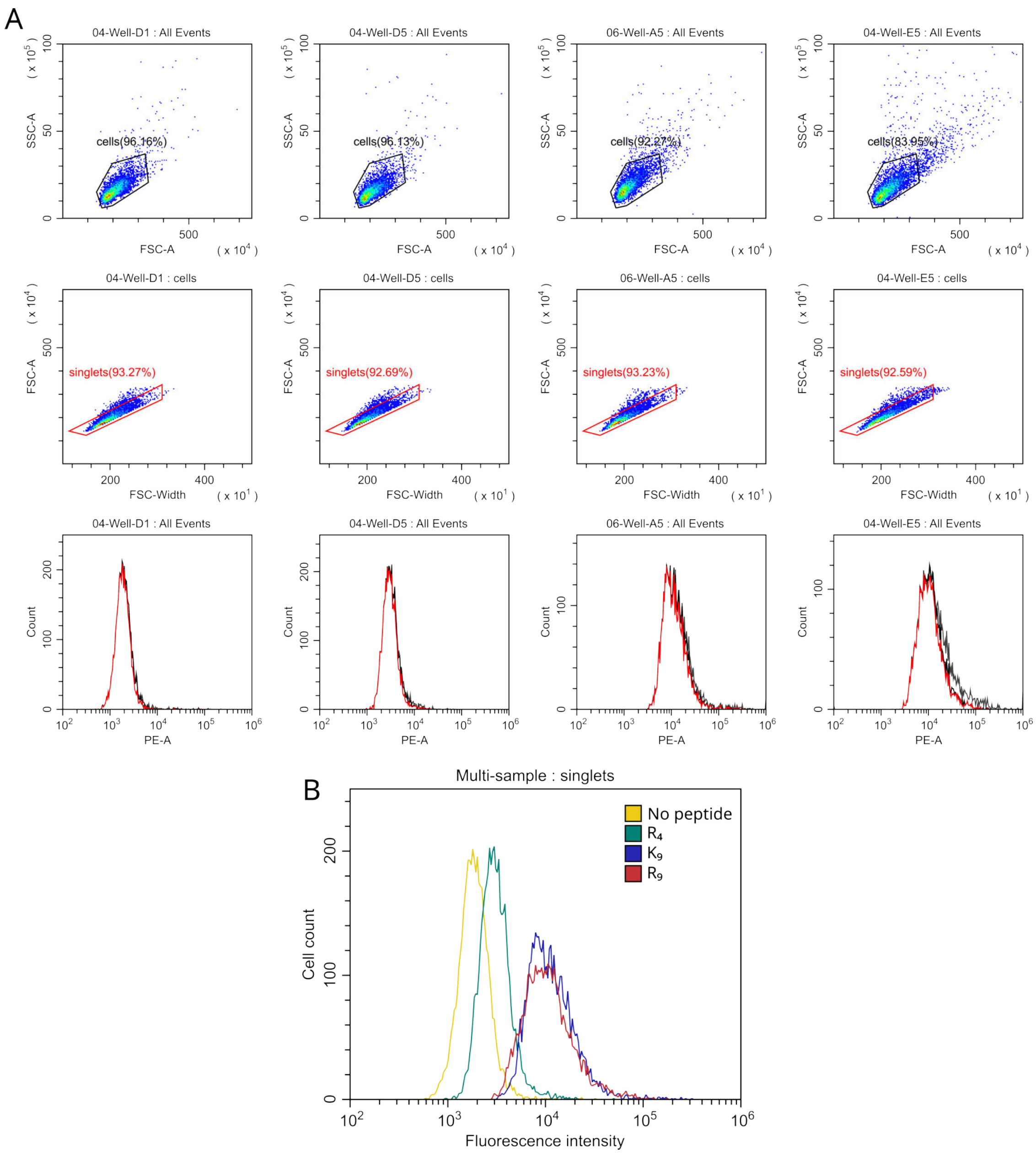


Figure S14: **Flow cytometry processing workflow example.** (A) First row shows live cell gating selection. In the second row, only singlet events are selected from the previous subpopulation. Third row shows resulting fluorescence intensity histograms for all detected cells (grey), "cells" gating (black), and singlets in red. Columns from left to right: cells without peptide, $R_4$, $K_9$, and $R_9$ (all 15 $\mu$M). (B) Resulting singlets histograms plotted together.

To compare, Figure S14, B shows the final fluorescence intensity histogram for the singlet cell populations incubated in HBSS buffer either without peptides (cells in yellow), with $R_4$ in green, $K_9$ in blue, and $R_9$ in red.

#### Time-Resolved Flow Cytometry

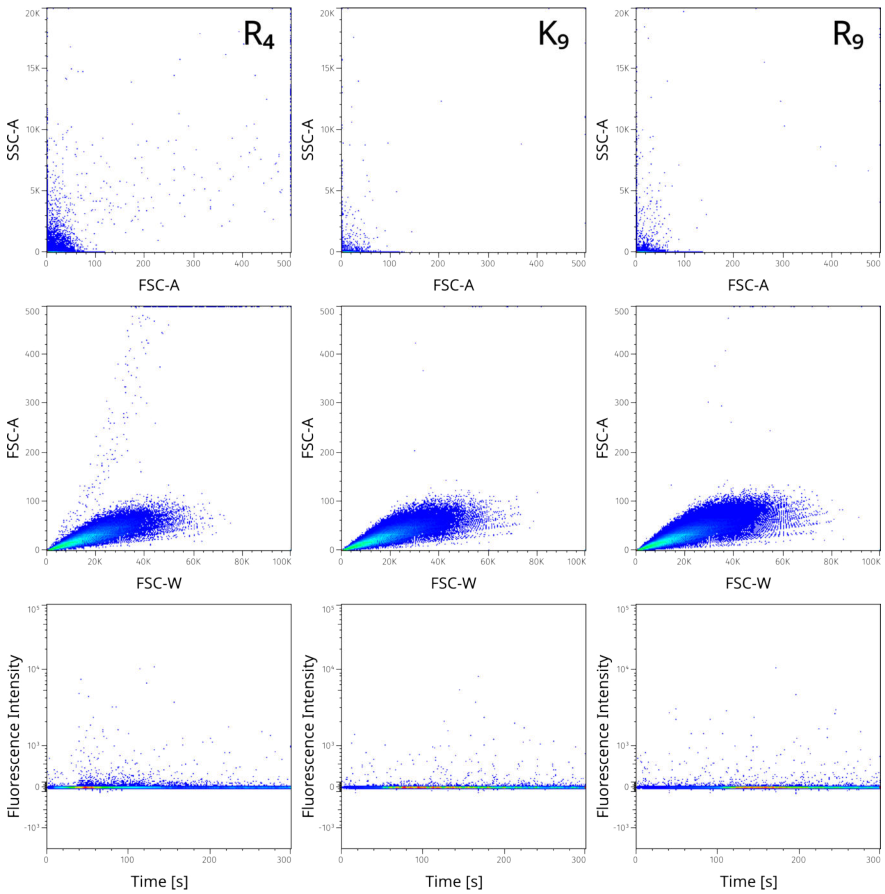


Figure S15: **Time-resolved flow cytometry** on a system containing peptides in HBSS buffer (without cells). Top row studies the size and internal complexity of the captured events. The middle row focuses on the size. The fluorescence time-evolution is shown in bottom row. Tetraarginine shown in left column, nonalysine in the middle, and nonaarginine is in the right column.

Without the presence of cells, Tamra-labelled peptides in the HBSS buffer (used for all cell experiments) were measured for 5 min by time-resolved flow cytometry, see Fig. S15. Smaller aggregates of various sizes are present (Fig. S15, top and middle rows). Similarly, no significant fluorescence intensity changes occurs in time (Fig. S15, bottom row).

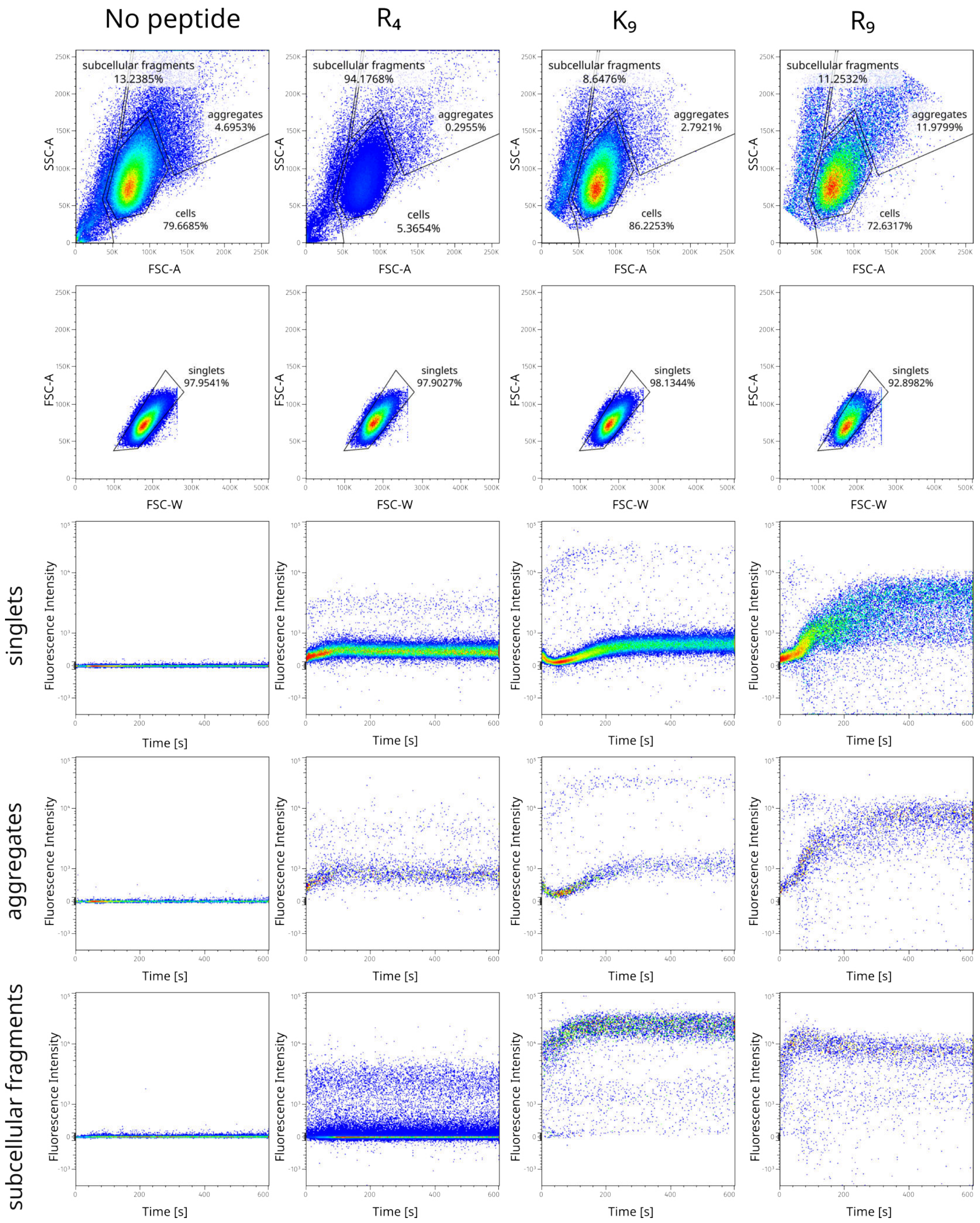


Figure S16: **Time-resolved flow cytometry processing workflow.** First row shows separating detected events into subcellular fragments (small), cells, and aggregates (large). In second row, singlet cells are selected from the cell population. The third, fourth, and fifth row illustrate peptide fluorescence intensity time measurements for singlets, aggregates, and subcellular fragments, respectively. Cells without peptide presence are shown in the left column, tetraarginine measurement is in the middle left column, nonalysine in the middle right, and nonaarginine is in the right column.

Concerning the time-resolved flow cytometry measurements, only the first 10 min of measurements was used for analyzis, since the fluorescent signal remained the same later. At the first gating step, subcellular fragments (small), cells, and aggregates (large) are selected, as shown in Fig. S16. Next, singlet events are extracted. Lastly, fluorescent intensity in time is plotted for all gated populations and different peptides.

### CLEM

Visualization of $R_9$-positive puncta on cell surface CLEM-based tomographic acquisition showed, alongside clear membrane remodelling, the presence of peptide aggregates and/or clusters which appear as dark amorphous blobs or puncta on the cryoEM grid (Fig. S17). The exact nature or composition of such clusters is currently unknown; the most likely scenarios could be aggregation of $R_9$ with other biomolecules, or the ultimate results of excessive membrane remodelling leading to lipid-peptide clusters.

In addition, we also observe high multilamellarity arising from vesicle-vesicle interaction (Fig. S18), similarly to the morphologies observed in cryoTEM.

While our tomographic acquisitions were performed on a single cryo-EM grid sample within one experimental session, we conducted a statistical analysis of the dataset to more rigorously assess whether the observed membrane-remodelling morphologies indeed represent a primary effect of the peptide's interaction with biological membranes. By analyzing the distribution of remodelling events across all visualized cells and the number of data points acquired per cell, we calculated the Wilson confidence intervals for the dataset and compared them with the global frequency of membrane remodelling (Fig. S19 A). Since the global remodelling probability lies within all confidence intervals, this provides statistical validation that the effect of $R_9$ is not dependent on the selection of a specific cell, but rather represents a general phenomenon.

Furthermore, because all observed membrane-remodelling events exhibited a multilamel-

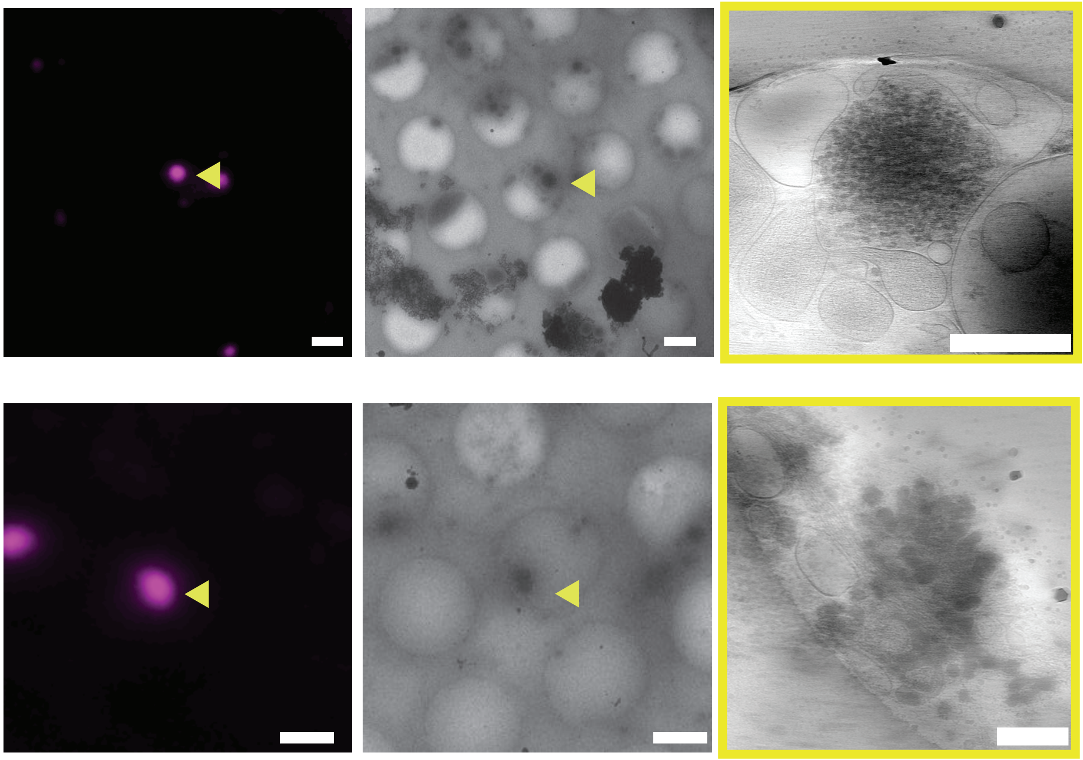

Figure S17: **Representative CLEM images of peptide aggregates or large peptide-positive clusters** resulting from incubation of U-2 OS cell sitting on an EM grid incubated with Tamra-labeled $R_9$, visualized both in cryo-fluorescence and cryo-electron microscopy (scale bar: 1 $\mu$m). Magenta represents Tamra fluorescence. High-magnification images show clear high electron density (scale bar: 200 nm)

lar morphology, we used statistical inference to estimate the probability of such an outcome given the overall frequency of multilamellarity as a remodelling phenotype (Fig. S19 B). The estimated p-value as a function of the multilamellarity probability indicated that, for the 27 observed events to all be multilamellar purely by chance, the global probability of an event being multilamellar among other possible remodelling morphologies would have to exceed 0.85. This strongly supports the notion that although other remodelling forms induced by the peptide cannot be excluded, multilamellarity constitutes the dominant outcome.

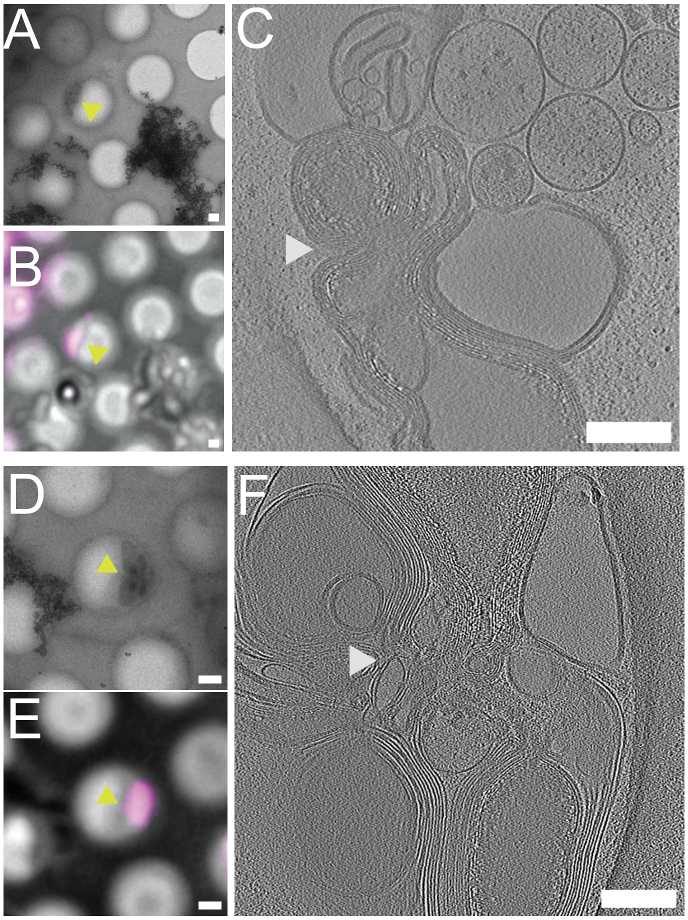


Figure S18: **Representative CLEM images of vesicle-vesicle interaction resulting in multilamellarity** Electron microscopy (A, D) and overlayed fluorescence and electron microscopy (B, E) images showing the point of interest with multiple vesicles (yellow triangles). Scale bar: 1 $\mu$m. (C, F) Corresponding high-resolution tomographic slice showing the high multilamellar morphology (white triangles). Scale bar 200 nm. Magenta represents Tamra fluorescence.

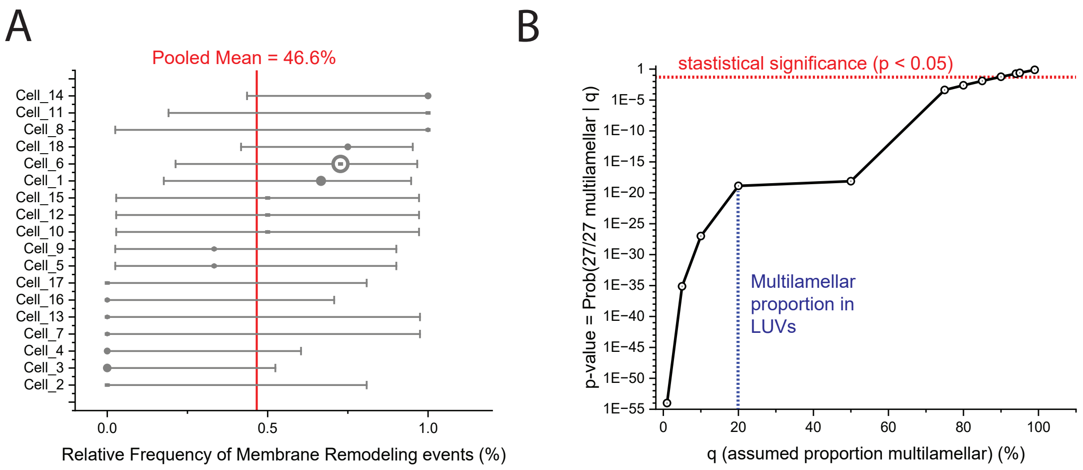


Figure S19: **Analysis of variability of events in CLEM datasets.** (A) Strip plot of multilamellar remodelling frequencies (%) per cell ($n = 58$ points, 27 remodelling events; global $p = 46.6\%$), sorted ascending. Point size scales with points imaged per cell ($n = 1$–$11$); error bars show 95% Wilson score CIs accounting for small sample sizes. All CIs overlap the global mean (horizontal line), forming characteristic Wilson funnel geometry—visual proof of sampling noise explaining 0–100% spread, not cell-specific effects. Non-significant Kruskal–Wallis test ($H = 17.00$, $p = 0.4544$) confirms uniformity. (B) Calculated p-values for observing 27/27 events of multilamellarity across various cells as a function of the probability of occurrence of multilamellarity.

(S3) Allen, M. P.; Tildesley, D. J. *Computer simulation of liquids*; Clarendon Press: USA, 1989.

(S4) Schachter, I. Lipid demixing reduces energy barriers for high-curvature vesicle budding. *Biophysical Journal* **2024**,

(S5) Helfrich, W. Elastic properties of lipid bilayers: theory and possible experiments. *Z. Naturforsch. C.* **1973**, *28*, 693–703.

(S6) Hamm, M.; Kozlov, M. M. Elastic energy of tilt and bending of fluid membranes. *Eur. Phys. J. E.* **2000**, *3*, 323–335.

(S7) Kozlovsky, Y.; Kozlov, M. M. Stalk model of membrane fusion: solution of energy crisis. *Biophys. J.* **2002**, *82*, 882–895.

(S8) Ryham, R. J.; Ward, M. A.; Cohen, F. S. Teardrop shapes minimize bending energy of fusion pores connecting planar bilayers. *Phys. Rev. E* **2013**, *88*, 062701.

(S9) Ryham, R. J.; Klotz, T. S.; Yao, L.; Cohen, F. S. Calculating transition energy barriers and characterizing activation states for steps of fusion. *Biophys. J.* **2016**, *110*, 1110–1124.

(S10) Ryham, R. J. Local changes in lipid composition to match membrane curvature. *Comput. Math. Biophys.* **2016**, *4*, –.

(S11) Wimley, W. C.; Thompson, T. Exchange and flip-flop of dimyristoyl phosphatidylcholine in liquid-crystalline, gel and two-component, two-phase large unilamellar vesicles. *Biochemistry* **1990**, *29*, 1296–1303.

(S12) Lipowsky, R. Multispherical shapes of vesicles highlight the curvature elasticity of biomembranes. *Adv. Colloid Interface Sci.* **2022**, *301*, 102613.

(S13) Mui, B.; Cullis, P.; Evans, E.; Madden, T. Osmotic properties of large unilamellar vesicles prepared by extrusion. *Biophys. J.* **1993**, *64*, 443–453.

(S14) Baumgart, T.; Hess, S. T.; Webb, W. W. Imaging coexisting fluid domains in biomembrane models coupling curvature and line tension. *Nature* **2003**, *425*, 821–824.

(S15) Jo, S.; Kim, T.; Iyer, V. G.; Im, W. CHARMM-GUI: A web-based graphical user interface for CHARMM. *Journal of Computational Chemistry* **2008**, *29*, 1859–1865.

(S16) Schachter, I.; Harries, D. Capturing Lipid Nanodisc Shape and Properties Using a Continuum Elastic Theory. *Journal of Chemical Theory and Computation* **2023**, *19*, 1360–1369.

(S17) Ayton, G.; Smondyrev, A. M.; Bardenhagen, S. G.; McMurtry, P.; Voth, G. A. Calculating the Bulk Modulus for a Lipid Bilayer with Nonequilibrium Molecular Dynamics Simulation. *Biophys. J.* **2002**, *82*, 1226–1238.
